# Putative condition-dependent viability selection in wild type stocks of *Drosophila pseudoobscura*

**DOI:** 10.1101/2020.04.10.036129

**Authors:** Ulku H. Altindag, Hannah N. Taylor, Chelsea Shoben, Keeley A. Pownall, Laurie S Stevison

## Abstract

Meiotic recombination rates vary in response to intrinsic and extrinsic factors. Recently, heat stress has been shown to reveal plasticity in recombination rates in *Drosophila pseudoobscura*. Here, a combination of molecular genotyping and X-linked recessive phenotypic markers were used to investigate differences in recombination rates due to either heat stress or advanced maternal age. However, haplotype frequencies deviated from equal proportions for crosses using phenotypic markers, indicating viability selection. Interestingly, skews in haplotype frequency were condition-dependent, consistent with the fixation of alleles in the wild type stocks used that are unfit at high temperature. Evidence of viability selection due to heat stress in the wild type haplotypes was most apparent on days 7-9 when more mutant non-crossover haplotypes were recovered in comparison to wild type (p=2.2e-4). Despite the condition-dependent mutational load in both wild type and mutant stocks, an analysis of recombination rate plasticity revealed days 7-9 (p=0.0085) and day 9 (p=0.037) to be significantly higher due to heat stress and days 1-3 as significantly higher due to maternal age (p=0.025). Still, to confirm these findings, SNP genotyping markers were used to further investigate recombination rate. This analysis supported days 9-10 as significantly different due to heat stress in two pairs of consecutive SNP markers (p=0.018; p=0.015), suggesting this time period as when recombination rate is most sensitive to heat stress. This peak timing for recombination plasticity is consistent with *D. melanogaster* based on comparison of similarly timed key meiotic events, enabling future mechanistic work of temperature stress on recombination rate.

## Introduction

Meiosis is fundamental for sexually reproducing organisms to generate haploid gametes. This process helps to maintain the correct number of chromosomes in the next generation, critical for gamete viability. Additionally, crossing over during meiosis creates novel genetic variation by recombining parental haplotypes, which can have important consequences for adaptation of species (Charlesworth & Barton, 1996; Page & Hawley, 2003).

Early studies in *Drosophila melanogaster* have shown that crossover rates vary as a result of various factors including maternal age, starvation, as well as external humidity and temperature (Plough 1917, 1921; Bridges 1927; Kohl and Singh 2018; Singh 2019). In more recent studies, it is shown that infection also alters recombination rate frequencies (Singh *et al*. 2015; Singh 2019). Over the last century, other model systems have replicated these results (reviewed in Parsons 1988; Agrawal *et al*. 2005; Bomblies *et al*. 2015; Modliszewski and Copenhaver 2017). For example, results from more recent studies indicate that desiccation is a recombinogenic factor and that desiccation-induced changes in both recombination rate and crossover interference are fitness-dependent, with a tendency of less fit individuals producing more variable progeny. Such dependence may play an important role in the regulation of genetic variation in populations experiencing environmental challenges (Aggarwal *et al*. 2019).

While these factors have consequences on events throughout meiosis such as in synaptonemal complex and double strand break (DSB) formation, early meiosis appears to be most sensitive to perturbation by a number of factors leading to apoptosis in these stages (reviewed in Stevison *et al*. 2017; Singh 2019). Experimental evidence points to temperature sensitive, pre-meiotic interphase as the stage when plasticity in recombination rate is the highest. This coincides with the relationship between meiotic recombination and DNA replication at S-phase (a component of interphase) (Grell 1973, 1978b).

While there has been a century of work on recombination rate plasticity in *D. melanogaster*, there have been no efforts to document this phenomenon in other *Drosophila* species. *Drosophila* is an extremely diverse genus made up of over 2000 species that diverged over 50 million years ago (Hales *et al*. 2015). Moreover, Parsons (1988) argued that *Drosophila* species can serve as indicators of global climate change due to their environmental sensitivity (Parsons 1988). However, one concern with focusing on *D. melanogaster* in the study of how environmental stress impacts recombination is that it is a cosmopolitan species, and thus may not have the same environmental sensitivity as other species within the *Drosophila* species group. Our team has recently worked to expand research on this ubiquitous phenomenon into *Drosophila pseudoobscura* (Stevison *et al*. 2017).

While traditionally studied for their inversion polymorphisms, *D. pseudoobscura* is native to western North America and a small region in Bogota, Colombia. It is therefore alpine over parts of its range, which means it has the potential to be more sensitive to environmental changes (Kuntz and Eisen 2014). This species of *Drosophila*, which is ∼30 million years diverged from the classic model, *D. melanogaster* (Throckmorton 1975), was the second *Drosophila* species to have its genome completely sequenced and is commonly used for chromosomal studies, which makes it a good model for recombination studies (Hales *et al*. 2015). Additionally, *D. pseudoobscura* females exhibit synchronization of oogenesis across egg chambers (Donald and Lamy 1938), which is key to studying the timing of events in meiosis because time is an indicator of progression through oocyte development. More recently, there has been a boost of interest in studying recombination rates in this species (Kulathinal *et al*. 2008, 2009; Stevison and Noor 2010; McGaugh *et al*. 2012; Samuk *et al*. 2020).

Our lab recently reported a preliminary analysis of recombination rate plasticity due to heat treatment during development in *D. pseudoobscura* (Stevison *et al*. 2017). In that study, significant plasticity was found in eight regions across the 2nd chromosome, with 5/8 regions showing higher recombination in the high temperature treatment (see Table S1 in Stevison et al 2017). These results parallel both classic and recent work done in *D. melanogaster* (Grell 1966, 1973, 1984; Singh *et al*. 2015; Ritz *et al*. 2017; Kohl and Singh 2018).

Here, this work was continued to establish *D. pseudoobscura* as a model for studying recombination rate plasticity. First, a series of experiments were conducted with the goal of pinpointing the timing of peak differences in recombination rate between control and temperature stress crosses. Temperature was used as treatment throughout development similar to work of Plough and others (Plough 1917, 1921; see figure 2 in Stevison *et al*. 2017), as well as maternal age. Phenotypic mutants were used and the experimental parameters were adjusted with each successive experiment, altering treatment between temperature and age, duration of progeny collection, progeny transfer frequencies, and sample sizes. Although the cross design primarily backcrossed to wild type flies to mitigate potential viability effects of the mutant markers, a thorough investigation into the haplotype frequencies from the mutant marker crosses was conducted to test for segregation bias. This analysis revealed these crosses to have significant deviations from the expectation of equal proportions based on Mendel’s first law. Interestingly, these results seemed to change between treatment and control as well as time points, suggesting condition-dependent variability in viability of the wild type alleles. Finally, SNP genotyping markers were used to confirm the recombination results from the phenotypic mutants due to their evidence for viability selection.

Combining strategies used in earlier studies, the work presented here provides important information for future mechanistic work to understand recombination rate plasticity and enable it to be studied in more depth in *D. pseudoobscura*.

## Materials and Methods

### Stocks

Genetic crosses using mutant markers were conducted using two X-linked recessive mutant *D. pseudoobscura* stocks. First, a double mutant stock was produced by crossing two lines obtained from the U.C. San Diego stock center (which has relocated to Cornell University): *yellow* (*y*; 1-74.5) found on the first chromosome (or chromosome X) at genetic map position 74.5 (stock 14044-0121.09, Dpse\y[1]) and *scarlet* (*st*) (stock 14011-0121.06, Dpse\v[1]). Mutations of the *scarlet* gene induce a bright red-eye phenotype (Beers 1937), and mutations within the *yellow* gene induce a yellow-hued body and wings (Sturtevant and Tan 1937). Second, a triple mutant stock (courtesy of Nitin Phadnis) was used that had three mutations - *yellow* (*y*; 1–74.5), *scalloped* (*sd*; 1–43.0), and *sepia* (*se*; 1–156.5) (Phadnis 2011). Mutations of the *scalloped* gene induce changes to the wing phenotype, whereas mutations in the *sepia* gene result in brown eyes (Crew and Lamy 1935). Genetic locations of all mutant markers are shown in Figure 1A. A fourth mutant in the triple mutant stock (*cut*; 1-22.5) produced inconsistent results likely due to a variation in penetrance of the mutation (Dworkin *et al*. 2009). Therefore, the *ct* marker was excluded from the remainder of the analysis.

**Figure 1.**
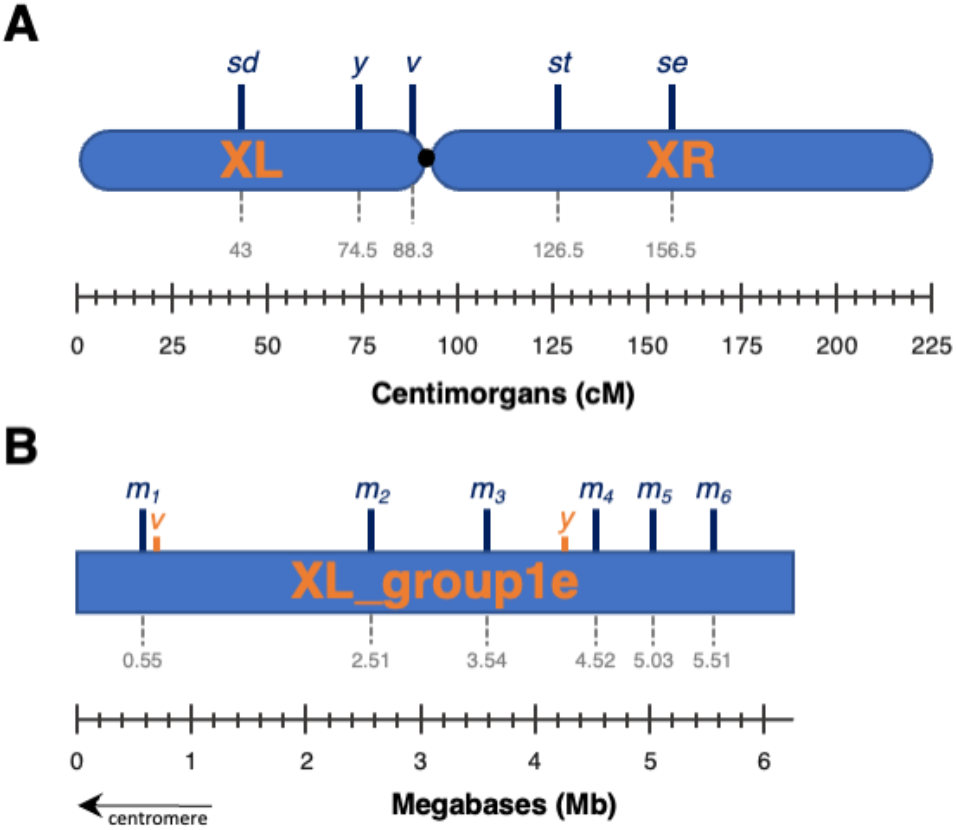
Physical and genetic location of markers used to measure recombination frequency in *D. pseudoobscura*. (A) Genetic map of X chromosome with location of mutant X-linked markers *scalloped* (*sd*), *yellow* (y), *scarlet* (st), and *sepia* (*se*) used to measure viability and recombination in Experiments 1-4. (B) Physical locations of SNP genotyping markers along 12.5 Mb scaffold “XL_group1e” located on the left arm of the X chromosome (XL). This scaffold (shown in reverse orientation) covers 62% of XL (only half shown here), including the mutant markers *vermillion* (*v*) and *yellow* (*y*).

Three wild-type *D. pseudoobscura* stocks were also used for genetic crosses. First, MV2-25 was used in crosses to the double mutant stock since it represents the reference genome strain (Richards 2005), and both are in an Arrowhead 3rd chromosome background. Second, to match the 3rd chromosome inversion arrangement of the multiple marker line, a second stock bearing the arrangement called “Treeline” was obtained from the National *Drosophila* species Stock Center at Cornell University (stock 14011-0121.265, Dpse\wild-type “TL”, SCI_12.2). This strain is also fully sequenced (NCBI Accession: SRX204785). Finally, AFC-57 (see Ritz *et al*. 2017)) was used for indel genotyping because it was a readily available wild type strain at the time.

### Husbandry & Cross Design

All stocks were maintained at 21°C with a 12 hour light-dark cycle in an incubator. Flies were reared on standard cornmeal–sugar–yeast–agar media in polypropylene vials.

For indel genotyping, all crosses were performed at 20°C in glass vials containing 6mL of corn syrup food. Virgin mutant female flies (5-7 days old) were crossed with male AFC-57. Virgin F_1_ females (5-7 days old) were collected and crossed with mutant male flies (Figure 2A). Resulting backcross progeny were phenotyped. Cross design for the SNP genotyping markers was described elsewhere (Stevison *et al*. 2017).

**Figure 2.**
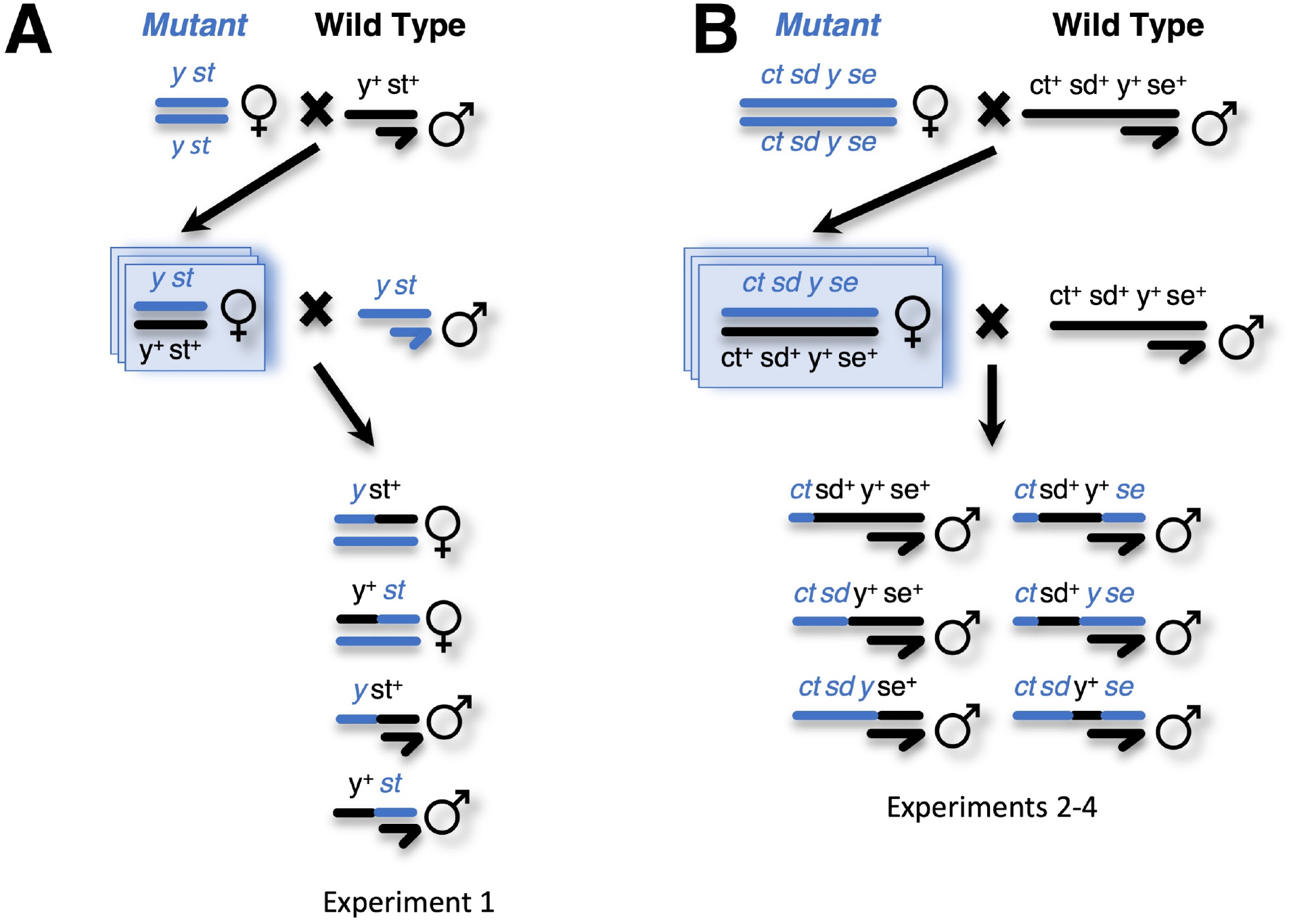
Crossing scheme for experiments using mutant phenotypes. Females homozygous for mutant markers of two stocks were used to cross to wild-type flies (indicated by plus sign). This F_1_ cross was the unit of replication, as indicated by the stacked boxes, and the resulting female progeny experienced the treatments as indicated in Table 1. The ID of these crosses were tracked in the resulting backcrosses. In Experiment 1, the *y-st* mutant stock and the MV2-25 wild-type stock were used (A). For Experiments 2-4, the triple mutant stock *sd-y-se* and the SCI_12.2 wild-type stock were used (B). Virgin F_1_ females were collected and stored in a common control temperature prior to the backcrosses. Based on initial screening of male backcross progeny, the marker *ct* was removed from consideration as it gave unreliable results due to incomplete penetrance. Male backcross progeny were screened for recombination analysis (Eq. 2) and female progeny were included for fecundity analysis (Eq. 1).

For genetic crosses, double and triple homozygous recessive mutant stock virgins were collected and aged 7 days to full sexual maturity. These flies were crossed to wild-type, age-matched males in control conditions to produce heterozygous F_1_ progeny (Table 1). Virgin heterozygous F_1_ females were collected within 8 hours of eclosion and stored at 21°C to maintain a common developmental timeline for treatment and control. There, they were aged to 7 days and backcrossed to wild-type males reared at 21°C. This cross design using wild-type males also provided a built-in ‘fail safe’ because female progeny could not be homozygous for the recessive mutant markers, and thus any mutant females would be an indicator of contamination.

**Table 1.** Summary of experimental design and results for genotyping experiments to measure the impact of temperature and age on recombination frequency. For each experiment, a different set of temperatures or ages, transfer frequencies and duration as well as sample sizes were used. *Note: Sample size is based only on the number of individuals targeted for recombination frequency. (e.g. only males were phenotyped for Exp 2-4). For fecundity, values are based on all progeny of both sexes for the duration of the experiment but not over the lifetime of each replicate female. For transfer frequency and duration, H denotes hours and D denotes days. ☥P-value from Eq. 1 for Treatment on Fecundity. ☨P-value from Eq. 2 for Treatment on Recombination Rate. Full anova tables for both analyses and *post hoc* tests are in Tables S2-S7. ^a^Crossing scheme matches Figure 2A. ^b^Crossing scheme matches Figure 2B. ^₺^Slashes in columns two through five indicate values split by treatment and control as indicated in the treatments column.

|  | Treatments<sup>‡</sup> | Transfer frequency/duration | # replicate crosses/treatment | Median # crosses/rep | Sample Size* | Fecundity<sup>¶</sup> | Recombination<sup>†</sup> |
| --- | --- | --- | --- | --- | --- | --- | --- |
| **Experiment 1<sup>a</sup>** | 20°C/25°C | 48H/6D;<br>72H/7-15D | 8/9 | 10/10 | 5187/4568 | 0.24 | 0.05 |
| **Experiment 2<sup>b</sup>** | 21°C/26°C | 72H/12D | 5/5 | 5/5 | 4140/2071 | **5.54E-11** | 0.76(sd-y),<br>0.91(y-se) |
| **Experiment 3<sup>b</sup>** | 21°C/26°C | 24H/6-10D | 6/6 | 12/12 | 3425/2084 | **4.49E-04** | 0.09(sd-y),<br>0.59(y-se) |
| **Experiment 4<sup>b</sup>** | 7-day/35-day | 72H/12D | 6/6 | 8/8 | 6219/4508 | **5.3E-3** | 0.93 (sd-y)<br>0.92 (y-se) |
| **SNP Genotyping** | 18°C/23°C | 48H/10D | 4/4 | 4/4 | 677/611 | n/a | 0.29 |

However, for Experiment 1, the backcross was done to the mutant stock (see below). Crossing schemes are diagrammed in Figure 2 with details on each experimental design outlined in Table 1, Figure 3, and below. Before backcrosses, wild-type males were individually isolated 24 hours prior to crosses to avoid crowding-induced courtship inhibition (Noor 1997). To backcross, a single wild-type male and single F_1_ female were placed in a fresh food vial. To increase sample sizes, multiple backcrosses were conducted from each replicate F_1_ cross using sibling female progeny.

**Figure 3.**
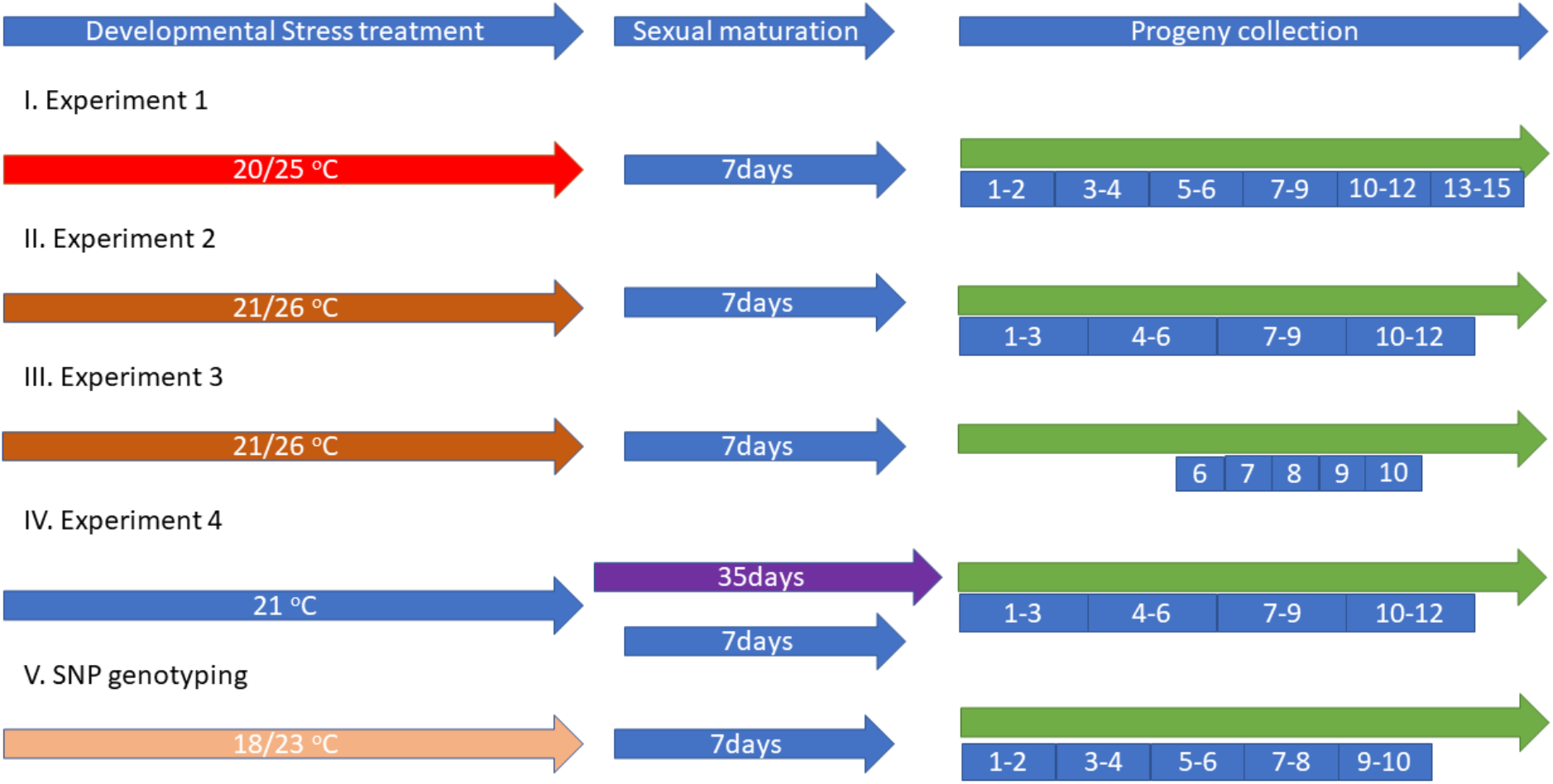
Experimental Design. Visual schematic of the experimental design for the series of experiments described. These parameters are also summarized in Table 1. For heat stress, F_1_ females experienced a developmental difference in rearing temperatures. For maternal age, females were either 7-days (control) or 35-days (treatment) at mating. For each experiment, mated F_1_ females were transferred with varying duration and frequency to partition the eggs laid into separate vials. All progeny from each vial as indicated by the blue boxes were collected for no more than a two-week period of time to avoid overlapping generations of progeny.

To promote mating, a cotton ball was placed inside to restrict available space and the vial was placed under a 100 Watt CFL light for an hour. After crosses, vials were assigned to identical incubators with a 12 hour light-dark cycle with the temperature varying according to Table 1 resulting in thermal stress throughout development. After 24 hours, the cotton was removed and the wild-type males were discarded to prevent additional stress from male harassment (Priest 2007). The female continued to be transferred to a fresh food vial according to the transfer frequency of each experiment (Table 1, Figure 3, and Table S1). Additionally, the vials where virgins were held prior to genetic crosses were kept for 14 days to ensure there were no larvae. If larvae were found, the cross was discarded.

#### Experimental Design

A series of four experiments were conducted using double (Experiments 1) and triple (Experiments 2-4) mutant stocks (summarized in Figure 3). First, an experiment was set up to investigate the impact of heat stress. The cross design for the first experiment was altered from the pilots to maximize sample size. Specifically, backcrosses were conducted to the X-linked recessive mutant stock rather than the wild-type stock as in the pilot experiments, allowing for the inclusion of female progeny in recombination calculations. Additionally, transfers were selected based on the aggregation of pilot experiment 1 data to hone in on the earlier time points with 48 hour transfers for the first 6 days and 72 hour transfers for the remaining 9 days, for 15 days total.

Next, to validate the findings in Experiment 1 using the triple mutant stock, Experiment 2 closely matched Experiment 1 modifying the transfer frequency to 72 hours for simplicity. Additionally, because there was no effect of temperature on fecundity in experiments at 25°C, the temperature treatment was increased to 26°C to increase the temperature stress. In Experiment 3, the 7-9 day post-mating time period was honed in with 24 hour transfers. However, to maximize the sample sizes in the later time points, both the number of replicates and crosses were increased relative to Experiment 1 and 2. Additionally, the vials where females were held for the first five days were discarded to keep the total sample size manageable.

Finally, to investigate the impact of maternal age, a fourth experiment was conducted closely matching the transfer frequency and duration of Experiment 2 (Figure S7). The heterozygous F_1_ females were aged to 7 days (control) and 35 days (maternal age treatment) and backcrossed to wild-type males. The collection, crossing, and F_1_ collection of the control flies were timed so they would be backcrossed at the same time as the 35-day old maternal age treatment flies. As shown in Figure S7B, the F_1_ females for the maternal age treatment were transferred into new vials every 7 days until they were 35 days old. When the maternal age treatment females were 35 days old and the control females were 7 days old, they were backcrossed to wild-type males.

SNP genotyping experimental design was described in (Stevison *et al*. 2017) and is summarized in Table 1 and Figure 3. The SNP marker design is described below.

#### Recombination Analysis Using Phenotypic Markers

Resultant progeny were screened for presence or absence of the mutant markers (Table 1). Except for Experiment 1, only male progeny were scored and if any female progeny were found to be mutant, the entire vial was discarded and the data removed. Visual scoring of mutant markers recorded each of the mutant traits independently in a single-blind manner. For Experiments 2-4, mutant scoring was delayed at least 5 days for the *sepia* eye color to become more pronounced.

Phenotyping ended 2 weeks after eclosion started to prevent the next generation from being included in the data. Data were entered in triplicate and compared until 100% concordant.

### Sequenom SNP Genotyping

As part of a preliminary characterization of plasticity in *D. pseudoobscura*, Sequenom SNP genotyping markers were designed to genotype crosses between FS14 and FS16 wild-type flies (methods previously described in Stevison *et al*. 2017; Sefick *et al*. 2018). Previously described results captured chromosome 2. In addition, for this study, six additional SNP markers were designed on the left arm of the X chromosome (chrXL) to span the region containing the mutant markers *yellow* and *vermillion* (Figure 1B). Together, the five intervals span 5Mb of the XL and are located on scaffold chrXL_group1e of the *D. pseudoobscura* reference genome.

### Molecular genotyping to investigate high recombination rate in double mutant

Molecular genotyping was used to confirm association between phenotypic mutants and their respective genes for the *yellow* and *vermillion* genes. For this analysis, two indel markers were designed based on the *D. pseudoobscura* assembly v3.1, each within 25kb from the *vermilion* and *yellow* genes. Markers selected resulted in differing PCR product length between the mutant stocks and the wild-type AFC-57 stock (Table S2). DNA was isolated (Gloor and Engels, 1992) from a minimum of 88 flies for each parent stock and backcross progeny for PCR amplification (Figure 2A). Length differences for markers were assayed via acrylamide gel. To confirm linkage between the *vermilion* and *yellow* genes and the red eye and yellow body phenotypes, backcross progeny of known phenotype were genotyped for the *vermilion-*linked and *yellow-*linked indel markers.

### Survivorship Analysis

In order to determine longevity of *D. pseudoobscura*, F_1_ females were generated using the same crossing scheme described for the recombination rate estimates.

Eighteen replicate crosses of 10 mutant females with 5 wild-type males were conducted, and the F_1_ female progeny were collected. Progeny were kept in vials with an average of 6.5 females (ranging from 1-13) based on when they were collected. To ensure the females had fresh food supply throughout the experiment, they were transferred to fresh food every 7 days. At each transfer, the number of females remaining in the vial was counted and recorded until no flies were left. For each replicate and time point, the percentage remaining as compared to the initial count was computed. The median across each time point was then computed to identify the time point at which less than 50% females remained. This analysis was used to justify the choice of age selected.

### Mutant Phenotype Segregation Analysis

Statistical analysis was performed using R v4.0.1 (R Core Team, 2020). For each experiment, haplotypes were grouped within crossover classes in order to investigate viability differences. The data from the backcross progeny were summed over up to 8 different types of haplotypes (Table 2). Additionally, the progeny were split based on both time point and treatment in Table 3. Because of the expectation of equal segregation of haplotypes during meiosis, a binomial test was performed in order to test for statistical deviations from 50-50 for each haplotype combination. Significant skews from expectation are indicated in bold with stars used to denote statistical significance (Tables 2-3). Additionally, the deviation from 50-50 was calculated across replicates for each CO class and treatment (Figure 4).

**Table 2:**
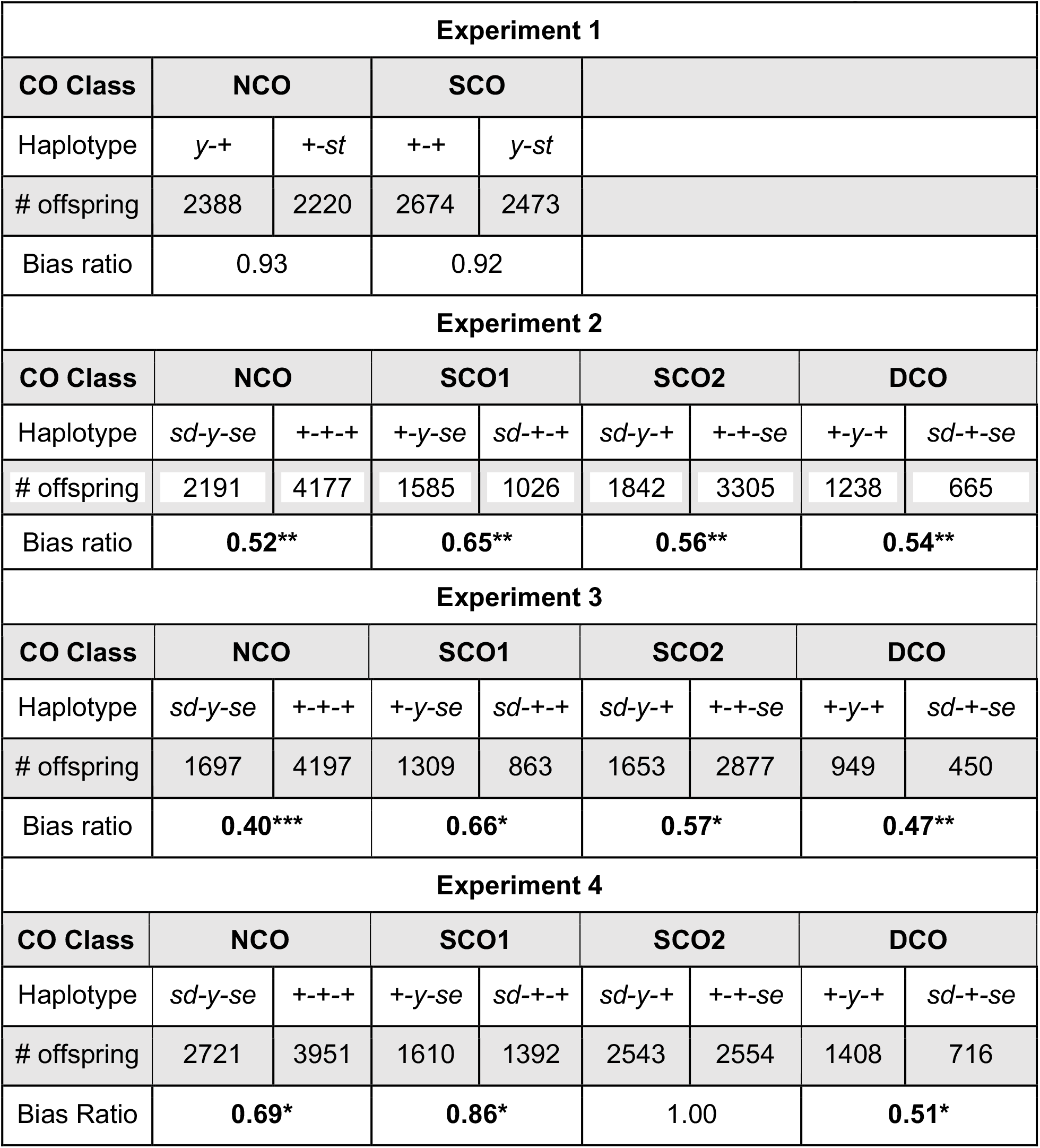
Haplotype frequencies for Experiments 1-4. Haplotype frequencies for each crossover class from each experiment using phenotypic mutant markers were analyzed to investigate possible segregation bias due to potential viability effects of visual markers for measuring recombination rate. A binomial test was performed to test for unequal proportions for each crossover class pair (bold values indicate significance; ***p-value<0.001, **p-value<0.01, *p-value<0.05). Bias ratio was calculated as the minimum divided by the maximum to keep values below 1 for better comparison. Additional breakdown of haplotypes by time point and treatment are shown in Table 3. Variation of bias ratio across F_1_ replicates and treatment are shown in Figure 4.

| Experiment 1 |  |  |  |  |  |  |  |  |
| --- | --- | --- | --- | --- | --- | --- | --- | --- |
| CO Class | NCO |  | SCO |  |  |  |  |  |
| Haplotype | *y-+* | *+ - st* | *+-+* | *y-st* |  |  |  |  |
| # offspring | 2388 | 2220 | 2674 | 2473 |  |  |  |  |
| Bias ratio | 0.93 |  | 0.92 |  |  |  |  |  |
| Experiment 2 |  |  |  |  |  |  |  |  |
| CO Class | NCO |  | SCO1 |  | SCO2 |  | DCO |  |
| Haplotype | *sd-y-se* | *+ - + - +* | *+ - y-se* | *sd - + - +* | *sd-y-+* | *+ - + - se* | *+ - y-+* | *sd - + - se* |
| # offspring | 2191 | 4177 | 1585 | 1026 | 1842 | 3305 | 1238 | 665 |
| Bias ratio | 0.52** |  | 0.65** |  | 0.56** |  | 0.54** |  |
| Experiment 3 |  |  |  |  |  |  |  |  |
| CO Class | NCO |  | SCO1 |  | SCO2 |  | DCO |  |
| Haplotype | *sd-y-se* | *+ - + - +* | *+ - y-se* | *sd - + - +* | *sd-y-+* | *+ - + - se* | *+ - y-+* | *sd - + - se* |
| # offspring | 1697 | 4197 | 1309 | 863 | 1653 | 2877 | 949 | 450 |
| Bias ratio | 0.40*** |  | 0.66* |  | 0.57* |  | 0.47** |  |
| Experiment 4 |  |  |  |  |  |  |  |  |
| CO Class | NCO |  | SCO1 |  | SCO2 |  | DCO |  |
| Haplotype | *sd-y-se* | *+ - + - +* | *+ - y-se* | *sd - + - +* | *sd-y-+* | *+ - + - se* | *+ - y-+* | *sd - + - se* |
| # offspring | 2721 | 3951 | 1610 | 1392 | 2543 | 2554 | 1408 | 716 |
| Bias Ratio | 0.69* |  | 0.86* |  | 1.00 |  | 0.51* |  |

**Table 3.**
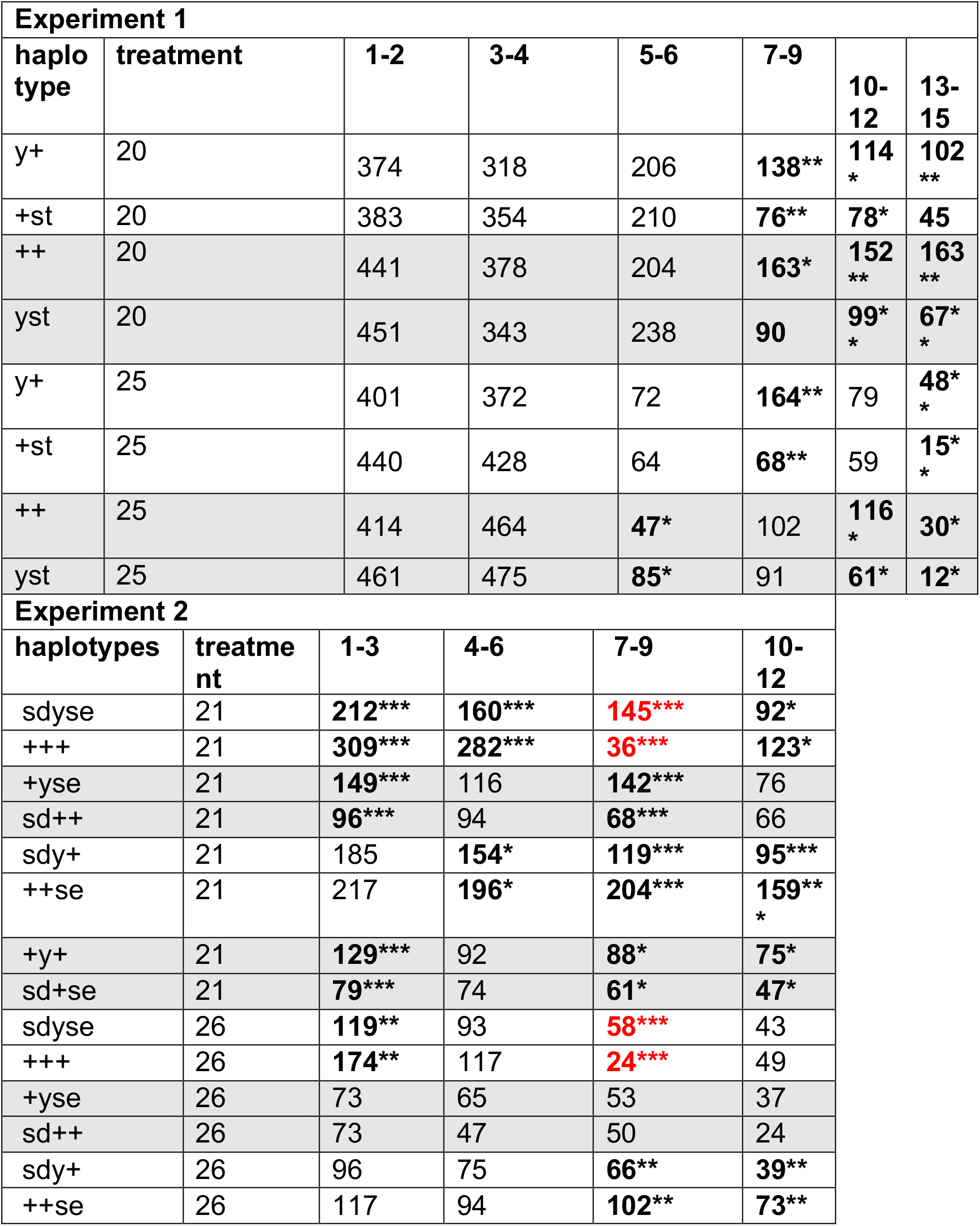
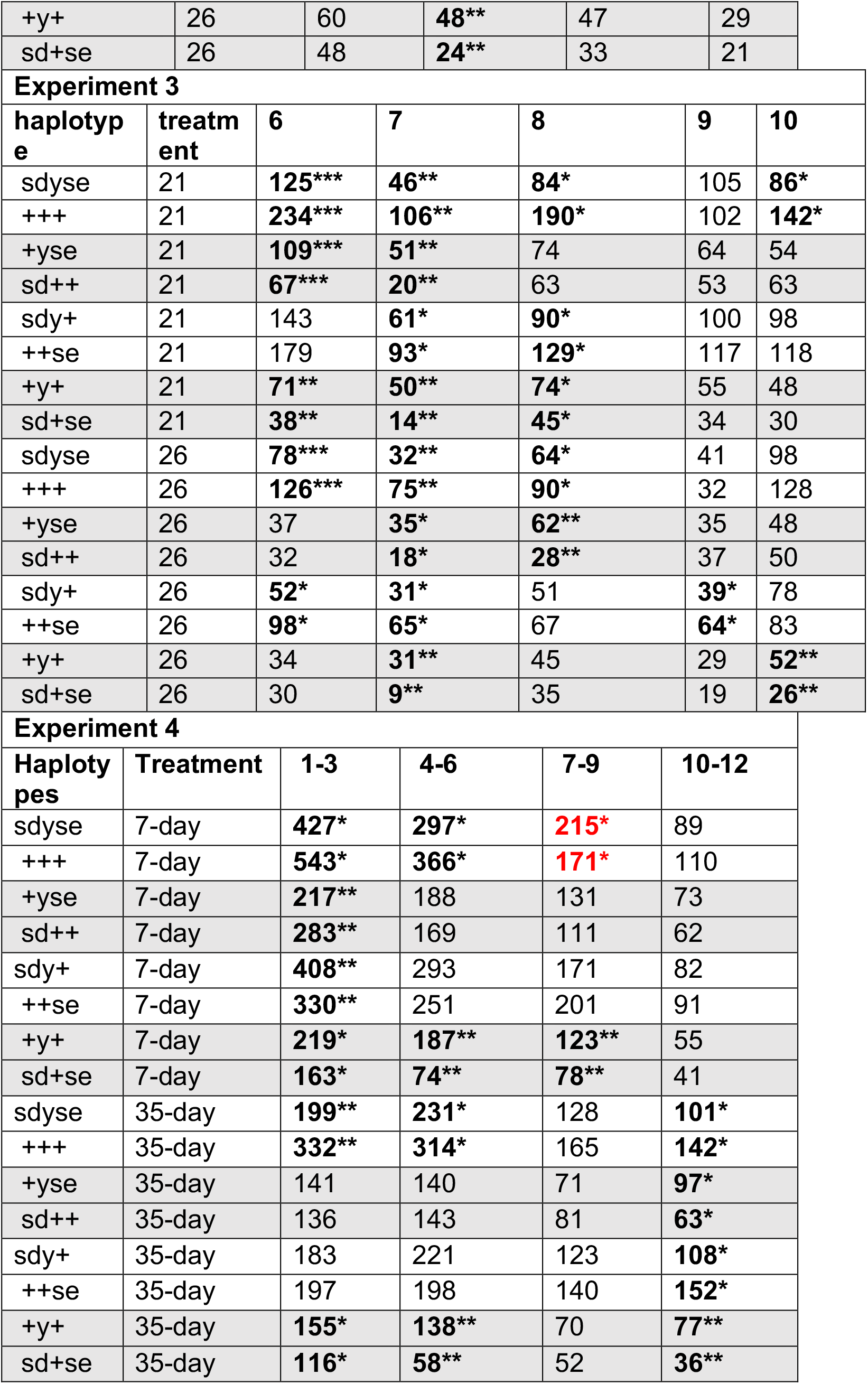
Haplotype frequencies for each experiment are provided as in Table 2, but here broken down further by treatment and time point. Binomial test was performed to test for the deviations between paired haplotype groups in the same experimental treatment/time point that should be in equal proportions (bold values indicate significance; ***p-value<0.001, **p-value<0.01, *p-value<0.05). Shown in red are NCO class crossovers where there is a significant excess of mutant progeny as compared to wild type.

**Figure 4.**
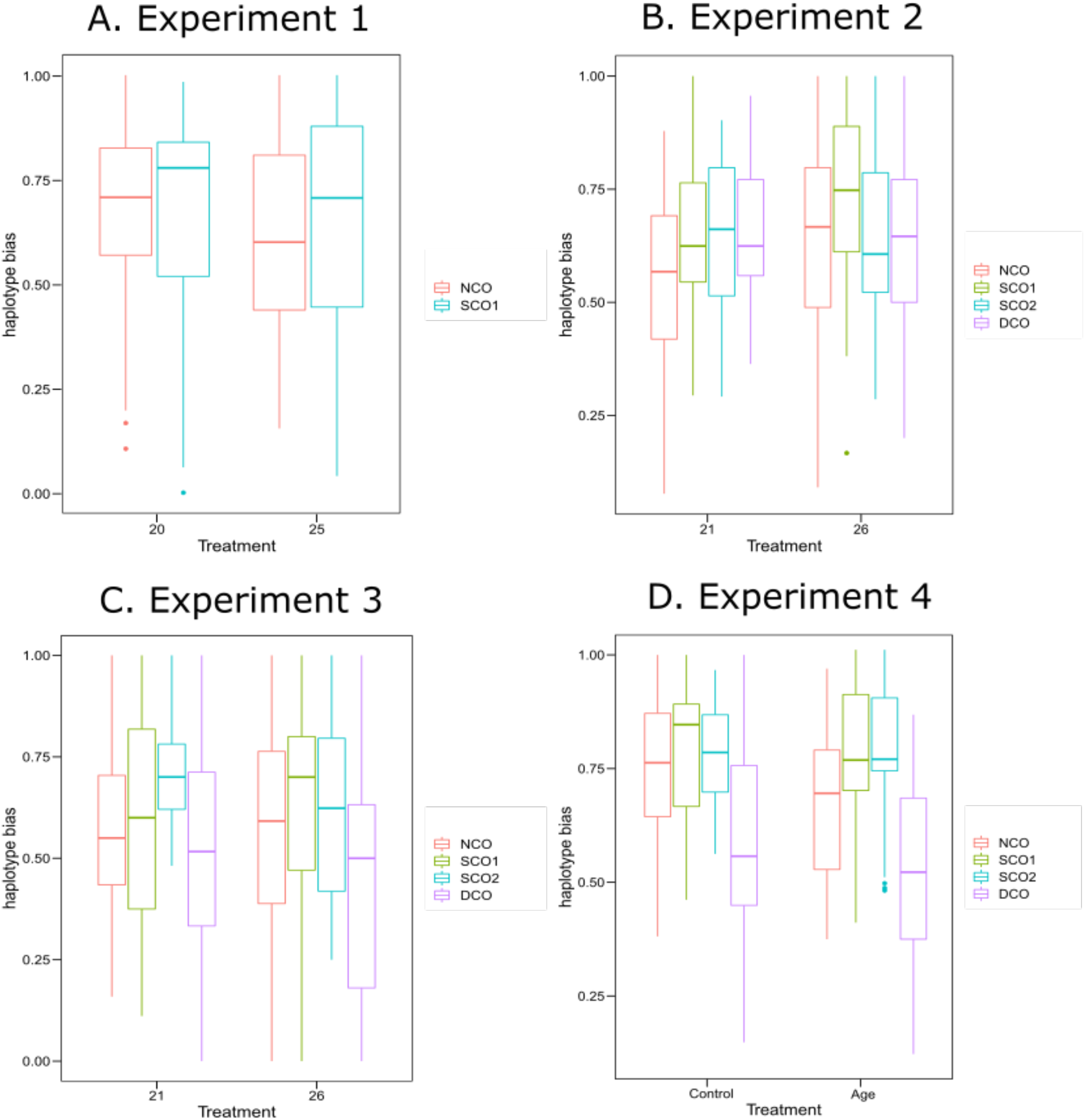
Condition-dependent viability results. Each panel features overall viability differences due to condition for each crossover class. Here haplotype bias was calculated by taking the ratio between the two haplotypes in the same CO class. For comparison, ratios were set up to always be below 1. Raw results are presented in Tables 2-3. Here, variability among F_1_ replicate crosses are shown. (A) For Experiment 1, due to having only two mutants, the NCO class has the largest difference in number of mutations per haplotype and the SCO class has an equal number of mutants between haplotypes. (B-C) For Experiments 2-3, which used a triple mutant stock, the SCO and DCO classes are both comparisons between one and two mutants. Whereas the NCO classes compare between three mutations and none. (D) Same as panels B-C, but for maternal age instead of heat stress.

### Statistical Analysis of Fecundity

Additionally, fecundity was tracked to measure the impact of stress due to temperature treatment and was calculated by dividing the number of backcross progeny to the number of F_1_ mothers. A quasipossion regression analysis was conducted following a similar basic model equation:

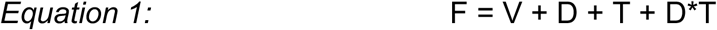

‘F’ indicates the continuous response variable of total number of progeny, or fecundity, for each time point. ‘V’ indicates the replicate vial ID and corresponds to F_1_ crosses. ‘D’ indicates the transfer period, or days post-mating, of the F_1_ female. Finally, ‘T’ indicates the temperature in which the F_1_ female was reared. For each replicate cross, fecundity was summed over all crosses and divided by the number of crosses per replicate to get an average number of progeny per time point for each replicate. Additionally, a *post hoc* lsmeans contrast was conducted to compute the significance of treatment versus control for each time point in each experiment (see Tables S7 and S8).

### Statistical Analysis of Recombination Frequency

Recombination rate frequencies were calculated for the chromosomal interval between each phenotypic marker (Figure 1A). Recombination frequencies (RF) correlating mapping distance between linked alleles were calculated by dividing the number of recombinant flies for regions *y-st, sd-y*, or *y-se* to the total number of progeny.

Glmer function was used to generate a fitted model using logistic regression per interval with replicate vial IDs as random effects and all other parameters as fixed effects. For each interval within each experiment, a logistic regression analysis with a mixed model was conducted in R. The basic model equation was:

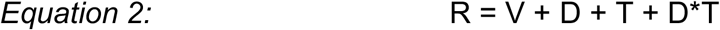

Here, all variables are the same as Eq. 1, except the response variable, ‘R’, in this model is the binary response variable of whether an individual offspring was recombinant or not based on the pair of mutant phenotypes over the screened region, for each time point. Progeny from backcrosses of F_1_ female siblings were summed per replicate cross per day and any replicate with fewer than 10 progeny were removed to avoid stochasticity in recombination rate estimates.

The results of both models are summarized in Table 1 and S1, and the full model tables can be found in Tables S3 and S4. Individual odds ratios were extracted for each time point using a *post hoc* means contrast between temperature and control to estimate biological relevance (Figures 5, S4 and S6). For logistic regression, exponentiating the coefficients of GLMM generates the odds of crossover formation between experimental and control conditions. A *post hoc* lsmeans contrast was done to calculate significance for each timepoint between treatment and control within the overall model for each experiment (see Tables S5 and S6).

**Figure 5.**
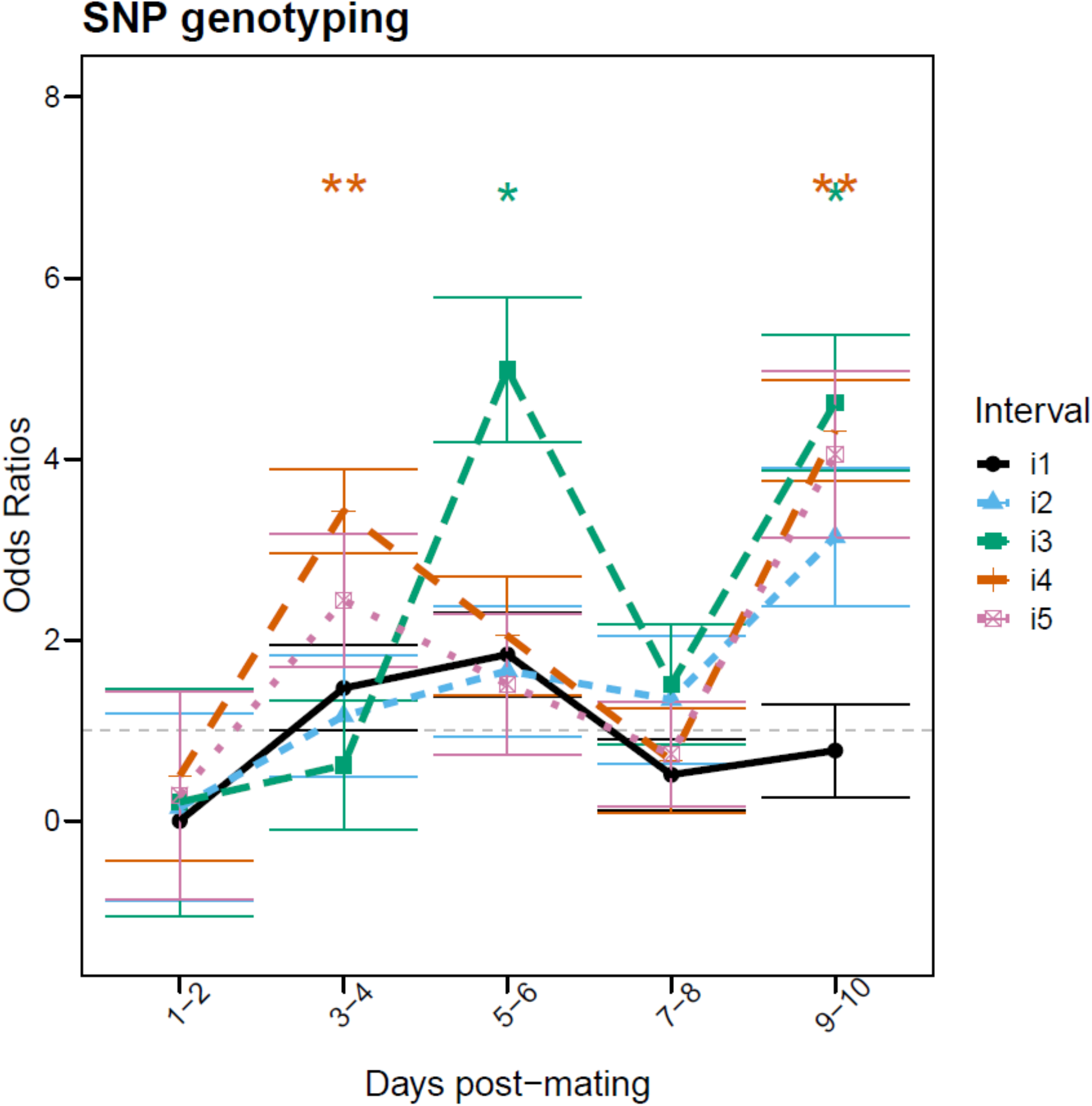
Recombination results for SNP genotyping recombination rate analysis. Recombination frequencies between control and treatment were compared using a fitted model using logistic regression. SNP genotyping markers span five intervals that overlap the *y-st* and *sd-y* intervals (Figure 1). In the overall model (Eq. 2) treatment was significant for intervals 3 (green) and 4 (orange) (Table S3). Exponentiating the coefficients generated the odds ratio. Odd ratios were plotted against days post-mating and indicate the odds of having a crossover in high temperature compared to control. A *post hoc* test was done to calculate significance for each timepoint between treatment and control with significance indicated via asterisks* (see Table S5). See Table 1 and Figure 3 for additional details regarding experimental design.

### Data Availability

Data files and scripts to complete the analysis are available on github at https://github.com/StevisonLab/Peak-Plasticity-Project. Zenodo was used to generate a DOI associated with a release of the code prior to publication: 10.5281/zenodo.XXXXXXX. This repository includes the raw survivorship data, raw mutant phenotype records for males, and female count data from the recombination analysis as csv files along with the R code. A separate csv file with treatment information includes dates and other metadata that would be needed to validate the analysis and conclusions herein. Additionally, a processed data file that includes sums of males and females, as well as crossover counts across each interval per time point, per replicate cross is also included. Also, a walk through tutorial for the analysis of data in Experiment 1 in Rmarkdown has been included. Sanger sequence data for 179bp of the *scarlet* gene, capturing the 2bp deletion, in the bright red eyed flies (originally ordered *vermilion* stock) is available on NCBI (GenBank accession number MT438819).

## Results

Experiments 1-4 used mutant markers which are known to have bias in haplotype frequency due to potential viability effects, therefore, we examined how this viability selection varied by treatment and time. We conducted a binomial test to determine if the differences in haplotype frequencies were significantly different from a 50-50 expectation (significant values bolded and starred Tables 2, 3, and S9). The four experiments showed condition-dependent variation in the overall skew from a 50-50 expectation (Figure 4).

### Double mutant cross reveals less overall viability selection

Experiments conducted using a double mutant stock crossed to the wild-type genome line MV2-25 (Figures 2A and S1) varied in transfer frequency and duration of progeny collection (Table 1 and S1). While the stock was labeled as *vermillion*, it was later discovered that the red eye mutant was associated with the *scarlet* gene rather than the *vermillion* gene (see Methods and Supplement), which explained differences in expected recombination frequencies in these experiments (see below).

Two smaller pilot experiments had smaller sample sizes than Experiment 1 (N=9,755, likely due to switching the cross design (Figure 2A vs. Figure S1). For the double mutant stock, the overall haplotype frequencies were not significantly different from equal proportions (Table 2). Unlike the overall data, there were some significant haplotype frequency skews that were most apparent at later time points and evident in both the control and high temperature crosses (Table 3). Specifically, there was a bigger skew in the two recombinant haplotypes, with the *y-+* haplotype being more frequent when frequencies were significantly different (Table 3). The most skewed proportions were found in the last time point on days 13-15 which had the fewest progeny. The next most skewed time point was the 7-9 day time period. The pilot crosses using this stock did not have nearly as much skew, which could be due to the difference in cross design (Figure 2A vs Figure S1). The majority of significant frequency differences in the pilot experiments were restricted to early time points and the control crosses (Table 2). In addition, because both males and females were phenotyped, haplotype frequencies were further examined for both sexes (Table 4 and S10). Unlike the non-significant variation between total progeny in Experiment 1, investigation based on sexes led to noticeable variation for both mutant and wildtype haplotype groups, but more skewed in female progeny (Table S10).

**Table 4.**
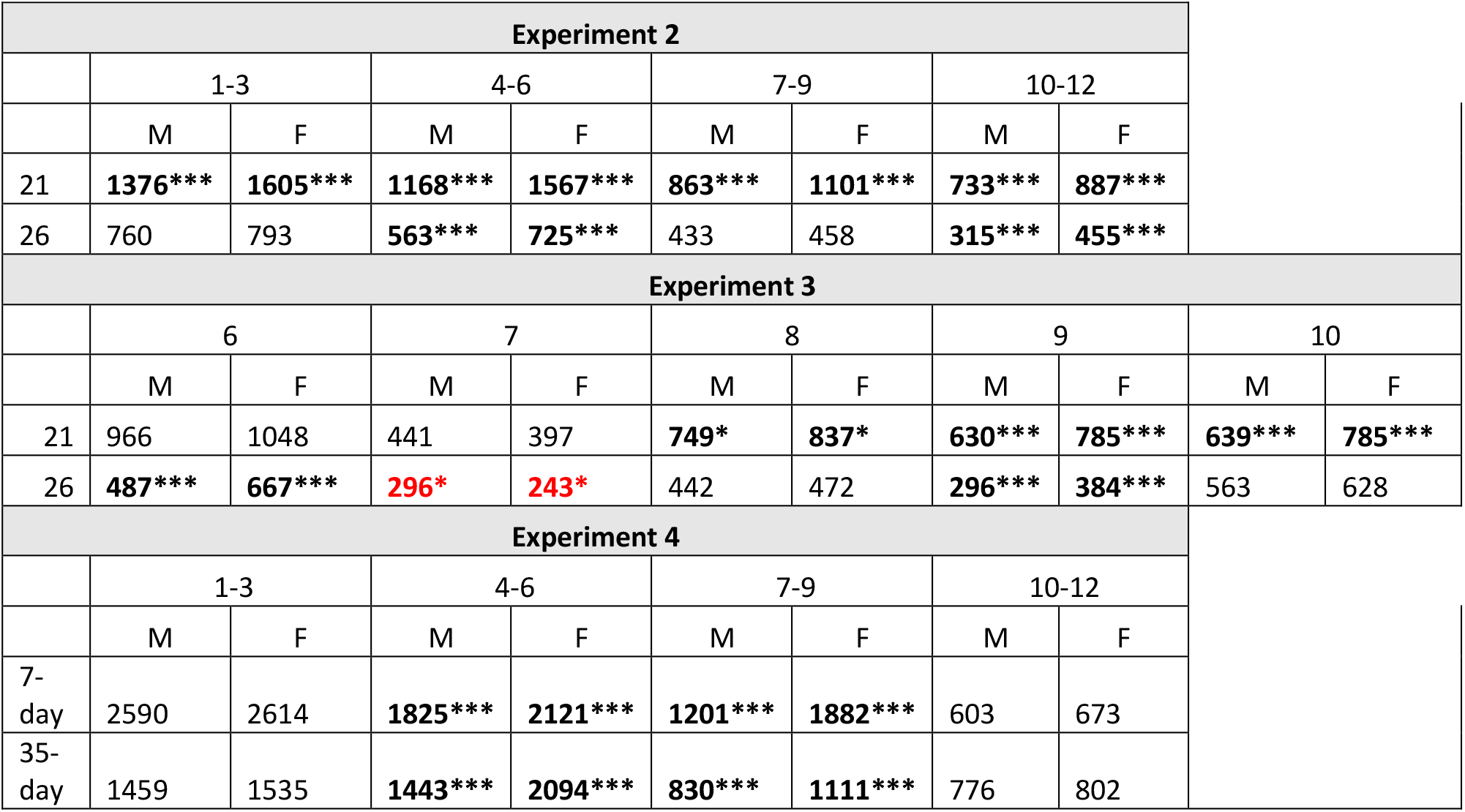
Male and female count data from mutant marker Experiments 2-4. Below, the number of male and female progeny per treatment and time period are shown. For Experiment 1, backcross was done to mutant stock (Figure 2A), so those results are split further by CO type in Table S10. However, for the other three experimental crosses, the backcross stock was wild type. Therefore, female progeny were never homozygous for the X-linked recessive markers and thus have no CO type information. Similar to the haplotype skew analysis in Tables 2-3, a binomial test was used to test for a significant deviation from 50:50 ratio (indicated in bold). Shown in red is the only case of a significant male bias in progeny.

### Triple mutant stock reveals strong condition-dependent viability selection

In Experiments 2-4, phenotypes at three mutant X-linked markers were recorded. For the triple mutant stock, the overall skew was much higher than in crosses with the double mutant stock (Table 2). Experiment 3 was most affected as a whole with a 2.47x difference in the proportion of NCO haplotypes (Table 2; p=0.0001) and 60% of haplotype pairs significantly different from equal proportions (Table 3). For recombinants, haplotypes with two mutant markers were typically lower in frequency than the alternative haplotype, with the exception being the *+-y-se* haplotype which is on average 1.41x higher than the *+-+-sd* haplotype (Table 2). This observation holds for all time points and treatments, with the exception being a 1.3x increase in the *sd-+-+* haplotype in days 1-3 for Experiment 4 (Table 3). This result suggests that the *scalloped* phenotype may contribute more to the bias in haplotype frequencies than the other mutant markers (but see below).

For Experiments 2-3, more than double the time points were significantly skewed in the control temperature as compared to the high temperature crosses, whereas for Experiments 1 and 4 both treatments had a similar number of skewed frequencies across time points (Table 3). Additionally, for Experiment 2 the 7-9 day time period had the most skewed haplotype frequencies. For Experiment 3, the 7 day time point had the most skewed proportions between haplotypes and the 9 day time point had the fewest skewed haplotype proportions. Finally, for Experiment 4, the 1-3 time point had the most skewed haplotype frequencies, predominantly in the control crosses; whereas the skew in haplotype frequencies in the day 10-12 time point are exclusively in the maternal age crosses (Table 3).

### Fecundity differences support stress of selected treatments

In Experiments 2 and 4, the selected treatment had a significant effect on fecundity (Table 1; Figure S2; Table S8), with a decrease in the treatment group indicating the stress response from the higher temperature of 26°C and the maternal age of 35 days. Similarly, fecundity declined steadily throughout progeny collection, consistent with a single mating event for these experiments. For Experiment 2, there was a 51% decrease in mean fecundity due to temperature (p<0.0001, see Table 1 and S8) that was significant for all time points (Figure S2). For Experiment 4, average fecundity for females aged 7 days used for the control crosses (70.36) differed from females aged to 35 days (54.29). A *post hoc* mean contrast found that fecundity was significantly different between treatments for the 1-3-day time point (p=1.46E-4) and the 7-9-day post-mating time point (p=0.013).

In Experiment 3, average sample size from days 6-10 in the control and experimental conditions were 20.9 and 15.0, respectively (p<0.019). Because the eggs laid by females on days 1-5 were discarded (Figure 3), this sample size does not represent lifetime fecundity. Still, the sample sizes were significantly different on days 6, 8, and 9 (Figure S2).

### Condition-dependent variation suggests viability selection of mutant and wild type alleles

When comparing the haplotype skew across time points and treatments, an interesting pattern emerges that sheds novel light on condition-dependent viability selection. For example, in Experiments 2-3, which had a significant overall reduction in sample size due to heat stress, the apparent skew is higher in control crosses as compared to high temperature crosses. One possible explanation is that the wild type stocks, being inbred laboratory strains held in a constant environment over many generations, have had fixation of alleles that are unfit at higher temperatures. This hypothesis is supported by the excess of mutant NCO class progeny in Experiments 2 and 4 seen in the 7-9 day time point (shown in red in Table 3). Assuming all mutant markers are equally unfit, the NCO class should show the largest skew against wild type as it has either three mutants or none. This switch in haplotype skew suggests that the wild type is also experiencing viability effects in addition to the visible mutant phenotypes for this treatment and time point. To further support this hypothesis, the skew is greater in control crosses for the NCO haplotypes than the heat stress crosses (Figure 4B). This is further supported by the above-mentioned skew in the SCO class where the *sd-+-+* haplotype has fewer progeny than the alternate haplotype which contains two mutant markers (*y* and *se*; SCO1 in Figure 4B). This skew is also significant for control crosses but not high temperature crosses in Experiment 2 for days 1-3 and 7-9 and Experiment 3 on day 6 (Table 3). A loss of wild type haplotypes at the higher temperature (due to homozygous wild type alleles that are temperature sensitive) could result in a reduced apparent skew in haplotype frequencies overall, leading to lower or no detectable bias in the high temperature treatment (Table 3; Figure 4). For Experiment 4, the bias in the crosses with increased maternal age do not see this reversal in the 35-day flies, suggesting it is specific to temperature stress. Therefore, the results suggests that the wild type stocks experience selection most at 26°C and 7-9 days post-mating. In Experiment 3, with 24 hour transfers, the NCO skew is significant for all time points except day 9 in both control and high temperature crosses, and day 10 for 26°C (Table 3). Similarly, the difference in NCO haplotype bias between temperatures is less apparent (Figure 4C), likely because it hones in on the time period 7-9 that is most skewed in Experiment 2. Together, these results suggest that mutational load of both mutant and wild type stocks are interacting to generate a condition-dependent pattern of haplotype bias.

To further investigate, the male-to-female ratios were evaluated (Tables 4 and S10). Based on the cross design which backcrossed to wild type males in Experiments 2-4, there is an expectation that the female progeny would exceed the male progeny if viability selection of the mutant markers were the reason for the haplotype skew. For Experiment 2 control, this is always true - males are significantly reduced as compared to females for all time points (Table 4). However, for 26°C, only time points 4-6 and 10-12 see significant female bias. Whereas time points 1-3 and 7-9 do not see any such bias. Similarly, for Experiment 3, there is a lack of female bias on days 8 and 10 at 26°C. For day 7, there is a significant excess of male progeny (p=0.025) at 26°C. This reduction of females as compared to males in 26°C crosses supports a viability effect of wild type alleles, consistent with the excess of mutant NCO progeny as compared to wild type NCO progeny on day 7-9 in 26°C reported above. This result supports the presence of alleles that are unfit at 26°C in the wild type stock. This phenomenon is largely absent from Experiment 4, where maternal age was varied instead of temperature. Specifically, time points 1-3 and 10-12 were lacking a female bias, but this was true for both the control and maternal age treatment, with no significant male bias.

Assuming this pattern is unique to the wild type stock used in Experiments 2-4, a similar analysis was conducted on the Experiment 1 data, where male and female progeny were analyzed separately. Interestingly, among female progeny, the 25°C crosses had more mutant than wild type NCO haplotypes on days 5-6 post-mating (shown in red in Table S10). For males, both treatment and control crosses had significantly reduced wild type NCO progeny on days 7-9 and 10-12; whereas for 25°C, the time point 3-4 is also significantly skewed against wild type progeny. This suggests the MV2-25 stock has similar fixation of alleles that are temperature sensitive, but at different time points and severity than the stock used in Experiments 2-4. Together, these findings suggest that the homogenous environment experienced by lab stocks fosters fixation of alleles that have lower viability across stressful environments (see Discussion).

### Recombination analysis of mutant markers inconclusive due to viability effects

Despite the condition-dependent viability found here, these experiments were further investigated for differences in recombination frequency over time and due to treatment. Of course, this was done with the understanding that when haplotype frequencies are skewed, an investigation of recombination frequency is flawed due to unrecovered haplotypes. Therefore, a genotyping experiment using SNP markers was used to confirm the differences in recombination rate along a similar region of the X-chromosome that the mutant phenotypic markers spanned (Figure 1B; see below).

For the double mutant stock, the recombination fraction calculated between the red eye and yellow body phenotypes were 46%, which prompted an investigation into the gene responsible for the red eye phenotype and ultimate discovery that it was due to a mutation in the *scarlet* gene (see Methods and Supplement). For Experiment 1, there was a significant interaction term between treatment and time point on recombination rate (see Eq. 2; p=0.0012, Table 1 and S4). Additionally, in a *post hoc* means contrast between treatment and control, days 7-9 post-mating had statistically significant higher odds of observing a crossover at high temperature (p=8.5e-2; N_20_=466; N_25_=424) with RF_25_=54.654% and RF_20_=50.54% (Figure 6A; Table S6). Additionally, days 13-15 were significantly different, but with a large standard error (see Table S6; Figure S6A) and small sample size with fewer replicates (N_20_=377; N_25_=97).

**Figure 6.**
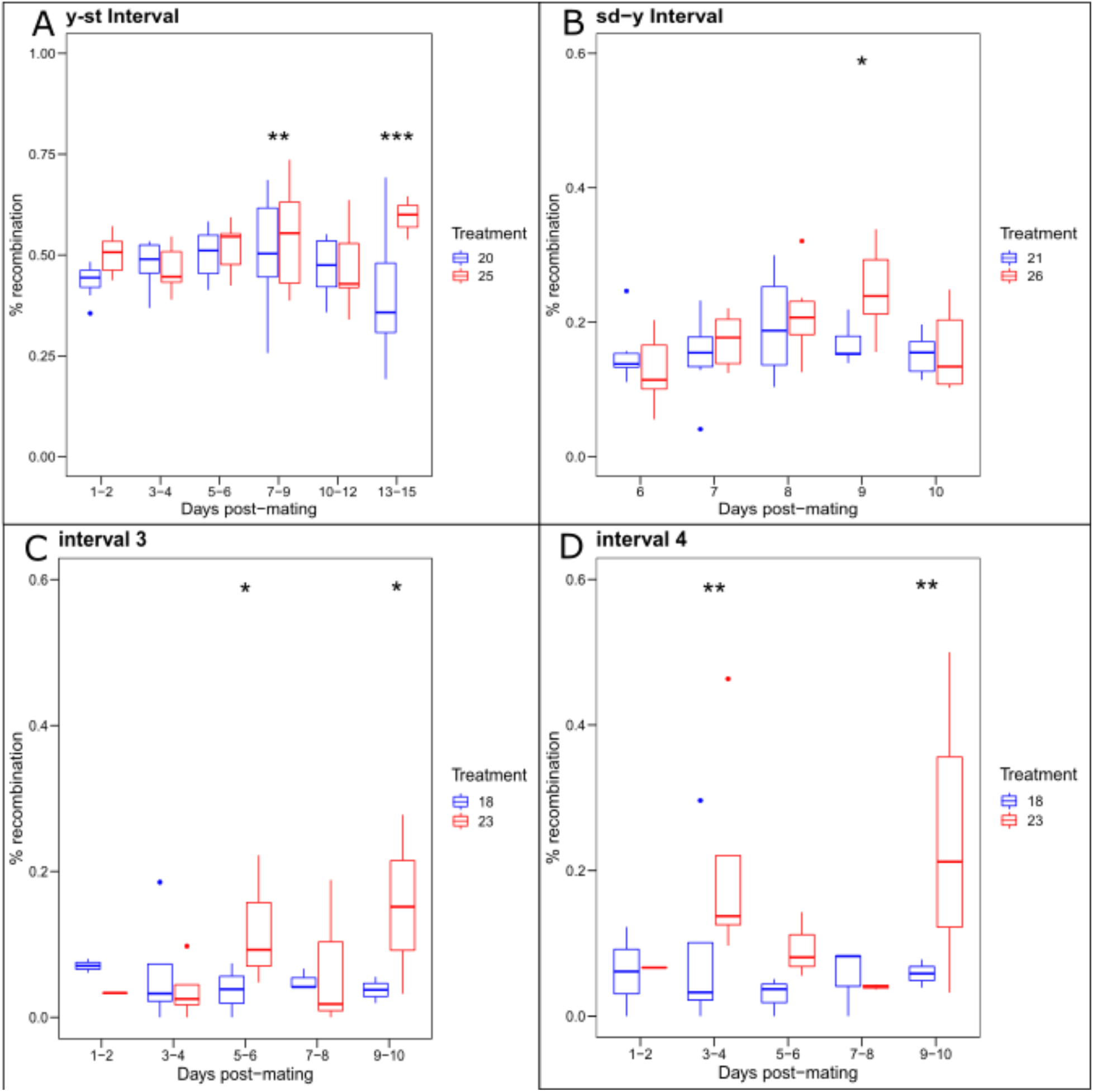
Recombination rate differences due to heat stress in mutant and SNP genotyping crosses. Each panel shows individual boxplots of the variation in the Kosambi corrected recombination rate among the individual F_1_ replicates per treatment. Significance between treatment and control for each time point in each plot are based on *post hoc* means contrasts and indicated by asterisks* (see Tables S5-S6). (A) As in Figure S6A, days 7-9 and 13-15 show significant difference in recombination rate between the control (blue) and high temperature (red) for the interval between *yellow-scarlet* from Experiment 1. (B) Similarly, the results from Figure S6C for the *scalloped-yellow* interval are shown from Experiment 3, where day 9 showed a significant difference in recombination rate. (C-D) Results from the SNP genotyping experiment for interval 3 (m_3_-m_4_) and interval 4 (m_4_-m_5_), which both show significant differences in recombination rate between control and high temperature treatment on days 9-10.

For the triple mutant stock, results for the *sd-y* region (32.1%) closely matched the expected rate (32.5%). Similar to *y-st*, the *y-se* region had a large recombination rate (46.0%) consistent with the genetic map distance (82cM), since markers over 50cM apart have a 50% recombination frequency. Kosambi corrections indicate lower recombination rates across both intervals (*sd-y*_kosambi_=20.1%; *y-se*_kosambi_=40%), perhaps due to the lack of recovery of all progeny as evidenced by the skewed haplotype analysis above.

For Experiments 2-4, the treatment did not significantly explain recombination rate in either interval for the overall model (Eq. 2; Table 1 and S4). A *post hoc* means contrast between treatment and control for Experiment 2 revealed that none of these results were statistically significant (Table S6; Figure S6B). In Experiment 3, the highest odds ratio was observed on day 9 (1.41) in the *sd-y* region (Figure S6C, p=0.034, N_21_= 630 and N_26_=296). Similarly, the median RF across replicates for this time point was 32.1% at 21°C and 40.4% at 26°C (Figure 6B). When the data for both intervals are combined, there was a 10.97% difference in total recombination on day 9.

In Experiment 4, although treatment was not significant in the overall model, the interaction between time points and treatment was significant (p=0.02; Table S4) for the *sd-y* interval. A *post hoc* mean contrast analysis revealed a significant difference in recombination rate (p=0.025; OR=1.16) in the first 72-hour time point for the *sd-y* interval (starred in Figure S6D; Table S6). Although heat stress and maternal age indicate different time points as sensitive to recombination plasticity, these results are inconclusive due to the extreme skews in recovered haplotypes noted above.

### Recombination rates for days 7-9 is reproducible between experiments

Above, recombination frequencies were inconclusive due to a lack of full recovery of progeny from viability selection of both mutant and wild type alleles. Therefore, it is worth determining how repeatable the measures are across experiments. However, due to the variation in experimental parameters (Table 1 and Figure 3), a direct comparison is not possible between all experiments. Still, the same time points from the same regions can be compared for a subset of experiments. Specifically, the 7-9 day post-mating time point can be compared between Experiments 2 and 3 *(sd-y; y-se*). The raw crossover count data from days 7-9 post-mating were aggregated from the 24-hour transfers in Experiment 3 into the same post-mating timeline of 72 hours in Experiment 2 for comparison. Odds ratios and standard errors were calculated and compared between individual experiment pairs (Figure S5 and S6). The results are highly similar with overlapping confidence intervals for both marker intervals (Figure S5).

### SNP genotyping markers confirm recombination plasticity of temperature sensitive time points

In an earlier molecular analysis, results were described for markers on the 2nd chromosome (Stevison *et al*. 2017). That analysis also included six X-chromosome SNP genotyping markers in the region spanning the genes *yellow* and *vermillion* on the X-chromosome (Figure 1B). In analyzing crossover data for intervals 1-3, the results show that control crosses had a 12.2% recombination rate, similar to the documented recombination fraction of 14.6 (Anderson 1993). The high temperature crosses had a 16% recombination rate across the same three intervals, which was significantly higher than the control (p=0.019).

Across the five intervals, a significant difference due to temperature was observed for interval 3, between markers m_3_ and m_4_, and interval 4, between markers m_4_ and m_5_ (Table S3). Additionally, a *post hoc* means contrast between treatment and control revealed a significant difference in recombination frequency (RF) in intervals 3 and 4 (Table S5). Specifically, interval 3 differed on days 5-6, and interval 4 differed on days 3-4. Both intervals 3-4 had a significant peak difference on days 9-10 (Figures 5 and 6C-D). Because intervals 3-4 overlap the *y-st* and *y-sd* regions, these SNP genotyping results are consistent with the sensitivity of recombination rate for similar time points (days 7-9 and day 9) and chromosomal regions as Experiments 1 and 3, respectively, described above that used mutant phenotypic markers. It is also worth noting that the magnitude of the difference due to temperature was higher for the SNP genotyping experiment than the experiments using phenotypic mutants (Figure 5 vs Figure S6).

## Discussion

Meiosis is taught in introductory genetics classes to be highly predictable and reliable, and yet for years scientists have been puzzled by deviations from the expectations set out by Mendel regarding the segregation of alleles. While many studies investigate haplotype skew, or transmission distortion, for evidence of unfit alleles (Meyer *et al*. 2012; Fu *et al*. 2020), the role of the environment to alter this skew is often ignored (but see Shoben and Noor 2020; Finnegan *et al*. 2021). Environmental heterogeneity is a known source of fitness differences and yet, the adherence to Mendel’s first law under various conditions has not been explicitly tested (Finnegan *et al*. 2021; Zwick *et al*.). Several studies have posited scenarios where competition among tetrads is variable across conditions suggesting recombination rate plasticity as a form of meiotic drive (Haig 2010; Stevison *et al*. 2017; Zwick *et al*.).

Biased haplotypes are a common observation when using mutant phenotypic markers, as certain genotypes are selected against due to viability effects, and are therefore not recovered in the progeny (Hurst 2019). Still, they offer an inexpensive alternative to test a variety of conditions and time points, which is why they were used here. While our investigation into haplotype frequencies complicated the initial purpose of our investigation, our data provided a unique opportunity to explore how different temperatures impact haplotype frequency and point to increased mutational load in wild type stocks. In this study, the segregation of the triple mutant gametes show the greatest skewed haplotype frequencies in the progeny, seemingly driven by the *scalloped* locus. However, a more thorough investigation into these results led to the conclusion that the wild type haplotype was being recovered with reduced frequency under high temperature stress across a select number of time points. Interestingly, this points to a mutational load in the wild type stock that is only revealed when reared at high temperatures. While the specific time points were not the same for the other wild type stock, similar results suggest this could be a more common phenomenon among laboratory stocks of Drosophila.

While it is certainly not unexpected for wild type stocks to harbor deleterious recessive alleles due to long term inbreeding, these are infrequently tested for such prior to their use in experiments. Moreover, for those that do investigate for the potential of viability selection in mutant or wild type stocks, this is likely only done in control conditions. Our results suggest that fecundity assays of wild type stocks should be conducted across a range of conditions before use in experiments. This is especially true for experiments that aim to investigate stress, meiotic drive, or recombination frequencies. In fact, our cross design is ideal for uncovering such condition-dependent viability selection in wild type stocks. For example, our design could be repeated with other wild type stocks to examine the variation in this phenomenon across stocks.

Further, our results suggest that fitness of lab stocks could be improved if they were reared under environmental heterogeneity to allow strains to purge unfit alleles that are sensitive across environments. This strategy should also be taken into consideration when establishing new lab stocks.

### Experiments point to days 9-10 as sensitive period for recombination rate plasticity

Similar to previous work (Stevison *et al*. 2017), we found a significant difference in recombination rate between flies reared at high temperatures as compared to control crosses for SNP markers on the X-chromosomes. However, only the model tables for SNP genotyping intervals 3-4 were significant for treatment, whereas the other experiments using mutant markers did not show a significant treatment effect. Further, *post hoc* analyses revealed various time points were significantly different between treatment and control with the most overlap between experiments on day 9 (9-10 in intervals 3-4 and 7-9 in Experiment 1; Tables S5-S6). The results from the experiments using phenotypic mutants were complicated by apparent viability selection in both wild type and mutant stocks, therefore, we focus our conclusions on the results from the SNP genotyping markers and heat stress. It is worth noting that the wild type stocks used for SNP genotyping were different than the ones used for the crosses with the phenotypic mutants.

A sensitive period of 9 days closely corresponds to work in *D. melanogaster* which suggests a similar sensitivity around day 6 due to heat stress. In *D. melanogaster*, development from oogenesis to egg maturation takes 10 days. Oocyte selection and development during oogenesis occurs in stages 1-14 in the last 79 hours (Koch and King 1966). Although, *D. pseudoobscura* oogenesis remains understudied, *Drosophila* species respond to temperature in a distinct manner. Still, a major benefit of *D. pseudoobscura* is the synchronicity of oogenesis among females that seems to alter with maternal age and indirectly affect fecundity (see Introduction). In *D. pseudoobscura*, eggs ripen as batches, with the immature eggs divided into groups of differing stages of development, ready to be deposited in large amounts at a time (Donald and Lamy 1938). Therefore, the number of eggs laid indicates a periodicity as compared to *D. melanogaster* that continuously lays their eggs in the 12 hours day/night cycles.

In a series of experiments, Grell was able to synchronize *D. melanogaster* eggs in age at the time of treatment, similar to the synchronicity observed in *D. pseudoobscura*. Her work identified variable expression of the gene *recombination defect (rec)* in temperature sensitive mutants of *D. melanogaster*. The protein encoded by the *rec* gene, MCM8, is involved in generating meiotic crossovers and DNA double strand break (DSB) formation has been shown to be evolutionarily conserved (Grell 1978a, 1984). MCM8 contributes to the stability of DNA strands during DSB and synaptonemal complex formation, and is transcribed early in developmental stages (Hunter 2015). In *Drosophila*, these events take place concurrently and affect regulation of crossovers (Carpenter 1975). The protein complexes common in these processes show a temporal pattern that can be tracked by developmental stages. Grell’s work in *D. melanogaster* showed that identifiable markers of DNA replication were present in 16-cell cyst in the adult flies by 6 days, pinpointing the peak plasticity at the same time. To identify the peak timing of recombination due to temperature stress in *D. melanogaster*, 6 hour transfers were conducted following perturbation (Grell 1973). While the experimental design in this study is quite different from Grell’s work, it is worth noting that in *D. pseudoobscura*, late replication domains indicated with markers of repressive histone marks and SUUR protein are present in the early stages of oogenesis, indicating the pre-meiotic S-phase occurs after day 8 post-mating coinciding with the observed peak in recombination rate plasticity in this study (Grell 1973; Higgins *et al*. 2012; Andreyenkova *et al*. 2013). In actuality, because females were held for 7 days to sexually mature, the peak corresponds to 15-16 days post eclosion.

These similarities between species suggests that the physiological processes influencing recombination rate need to be further explored in a comparative context. Although there has been a lot of work done in *D. melanogaster*, there are other *Drosophila* species that may be more sensitive to environmental perturbations for studying this important phenomenon. Here, we have examined plasticity in the alpine species, *D. pseudoobscura*. Additionally, cactophilic (Markow 2019) and mushroom feeding (Scott Chialvo *et al*. 2019) *Drosophila* represent recent adaptive radiations with growing potential for ecological genomics. Finally, the *montium* species group has recently become genome-enabled and is well suited for testing various evolutionary hypotheses (Bronski *et al*. 2020).

## Supporting information

Supplementary Methods and Results

## Acknowledgements

This work was supported by research start-up funds to LSS from the Department of Biological Sciences at Auburn University. UHA was supported by NSF-DEB EAGER No. 1939090 to LSS. We thank members of the Noor Lab for generating the double mutant stock used for Experiment 1. We thank Mohamed Noor for guidance and feedback on this work. We thank the Graze lab for help with fly husbandry for pilot experiments and Experiment 1, which were conducted in the fly lab of Rita Graze of Auburn University. We thank the Phadnis Lab for sharing the triple mutant stock used for Experiments 2-4. We thank Todd Steury for consultation on the statistical analysis. We thank members of the Stevison Lab for extensive help with recording mutant marker phenotypes, with particular thanks to Natalia Rivera-Rincon, Neeve Curley, Kyle Meding, Anna Tourne, Adam King, Kaitlyn Walter, and several other undergraduate researchers over a three year period. This work is part of the USDA NIFA Hatch project ALA0021-1-18015.

## article summary

Meiotic recombination contributes to genetic variation by creating different allelic combinations and is often deemed beneficial for eukaryotic organisms. Factors such as temperature and age induce recombination rate plasticity. Even though plasticity in *Drosophila* has been studied extensively, this work has been primarily done in *D. melanogaster*. Mutant phenotypic markers present rapid and inexpensive options for studying this phenomenon, yet are subject to viability selection. Here, condition-dependent viability selection results suggest wild type *D. pseudoobscura* stocks similarly experience viability selection under stress. Still, SNP genotyping markers revealed the peak timing of recombination rate plasticity to occur around 9-10 days post-mating.

