## Supplementary Methods and Results for "Putative condition-dependent viability selection in wild type stocks of *Drosophila pseudoobscura*"

### Pilot Recombination Rate Experiments using double mutant stock

Based on the experimental design in the SNP genotyping experiment, two pilot experiments were conducted using the mutant stock *y-st*. First, the duration of progeny collection was extended from 10 days to 15 days. Additionally, the transfer frequency was increased from 48 to 24 hour transfers. The control temperature of 18°C was maintained, but the temperature treatment was increased from 23°C to 24.5°C to maximize differences between the treatment and control (Table S1), careful to avoid going beyond upper bounds of developmental temperatures reported in the literature as 25°C (Kuntz and Eisen 2014)

For this first pilot experiment, several time points had a low sample size with too few replicates and missing data for either treatment or control. Therefore, time points were aggregated for analysis (Figure S4A). Still, recombination differences when aggregating all time points ($RF_{18}$=44.3%; $RF_{24.5}$=46.7%) were not statistically different.

A second pilot experiment, based on the differences at the later time points of days 10-12 and 13-15 in the first experiment (see results), the duration was extended even further from 15 to 20 days, but the transfer frequency was considerably widened from 24 hours transfers to 120 hour transfers (Brenman 2017). The control temperature was shifted from 18°C to the more standard rearing environment of 20°C (Smith 1958). To maintain a 5°C difference between treatment and control, the temperature treatment was also shifted slightly to 25°C (Table S1).

Still, the second pilot failed to show any statistically significant differences in RF in any individual time points (Figure S4B), or when aggregated ($RF_{20}$=44.7%; $RF_{25}$=47.8%). Additionally, due to the experiments having only a single mating event, the progeny sample sizes for days 16-20 in the second pilot were too small for further consideration.

Rearing temperature did not significantly explain differences in fecundity (Eq. 1) across either pilot experiment (Tables S1, S7, and S8). However, the first pilot showed a decrease in the mean fecundity between temperatures of 17% (Figure S2A).

Both pilot experiments showed a higher RF in the temperature treatment than the control (Figure S4A-B; n.s.), with the second pilot having a slightly larger effect size, likely due to the adjustment to temperature. Although there was not enough power to say whether these were significant differences, likely due to low sample size (Table S1), they were useful in guiding the design of Experiment 1 described in the main paper.

In the pilot experiments, the recombination fraction calculated between the red eye and yellow body phenotypes were 46%, much higher than the published 14.6%, indicating that one of the mutants was not in the expected gene region. For the indel marker near the *yellow* gene, results indicated an association between genotype and phenotype (recombination fraction 0/88). However, upon genotyping an indel marker near the *vermillion* gene, it was discovered that the bright red eye phenotype did not associate with the indel marker among backcross progeny (recombination fraction 39/85).

### Molecular genotyping reveals red eyed stock is actually *scarlet*

Next, a mutation in *scarlet (st*), another X-linked marker in *D. pseudoobscura*, which also causes a red-eye phenotype was investigated. Genotyping confirmed that the bright red eye phenotype was 100% associated with an indel within 21.9kb of the *(st)* gene (recombination fraction 0/471). Sanger sequence analysis revealed a 2bp deletion within the *scarlet* gene of the bright red eyed flies of this strain, suggesting a frame-shift mutation within the *scarlet* gene is a likely cause for this phenotype. Although there is not a consistent map location for *scarlet* in the literature in *D. pseudoobscura* (Beers 1937; Mampell 1943), it is consistently 30cM away from *sepia* (Figure 1B). This places it roughly 52cM away from *yellow*, consistent with the observed recombination rates in both the pilot experiments and Experiment 1 here.

### Median survivorship is 42 days of age

To investigate the impact of maternal age on recombination rate, the longevity of this species was followed to select an appropriate age. For this analysis, 808 $F_1$ virgin female progeny across 18 replicates were followed until none remained (91 days). Each replicate cross had an average of 46.28 females ranging from 8 to 75 (housed in smaller groups, see Methods). Holding the females as virgins could have an impact on longevity, as females in the wild mate multiples times in their lifespan. The percentage of survival decreased slowly between day 0 and day 30, rapidly between day 30 and day 57, and slowly between day 57 and day 91 (Figure S3). The inflection point of the survivorship curve where the median survivorship was below 50% occurred at 42 days. At 42 days, 61% of replicates had more than 50% loss as compared to original counts. The time point immediately preceding this was 35 days. At 35 days, only 22% of replicates had greater than 50% loss. Therefore, 35 days was selected as the treatment for the mutant screen experiment to investigate how maternal age impacts recombination rate.

### Fecundity and survivorship results support reduced fitness due to maternal age

Reduced natural selection later in life has been implicated for age-related fitness declines, such as fecundity and longevity. The fecundity and survivorship results here were consistent with decreased fitness with maternal age. Fertility of *Drosophila* depends on successful oogenesis, ability to store sperm, fertilization success, and the age of the parents. In *D. melanogaster*, oogenesis is affected by maternal age and the number of oocytes produced decreases as the age of the mother increases (Brenman 2017). Additionally, a 50% decline in fecundity is observed between mothers aged 22-28 days old and mothers aged 4-7 days old in *D. melanogaster* (Miller *et al.* 2014). In the survivorship analysis here, females lived up to 91 days, and it was not until 42 days of age that longevity reduced below 50%. Fecundity in Experiment 4 showed the expected significant decrease (22.84%) between 35-day old and 7-day old females (Figure S2F). Additionally, the survivorship results here (Figure S3) were consistent with a type 2 survivorship curve, which represents increased variation in age at death among individuals, as compared to type 1 and type 3 curves (Demetrius 1978).

### Supplementary Figures

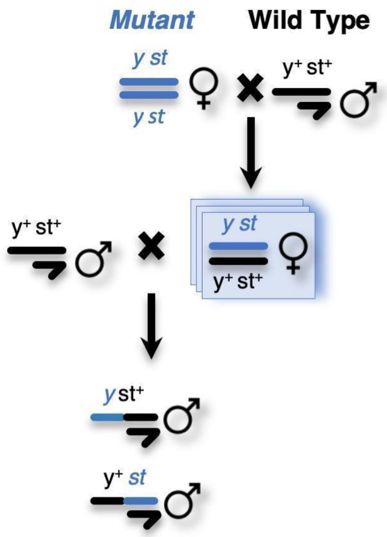


**Supplementary Figure 1. Crossing scheme for pilot mutant phenotype screens using double mutant stock.** Females from the *y-st* mutant stock were used to cross to males from the MV2-25 wild-type stock. This $F_1$ cross was the unit of replication, as indicated by the stacked boxes, and the resulting female progeny experienced the developmental difference in rearing temperatures as indicated in Table S1. The ID of these crosses were tracked in the resulting backcrosses. Virgin $F_1$ females were collected and stored in a common control temperature prior to the backcrosses. Male backcross progeny were screened for recombination analysis (Eq. 2) and female progeny were included for fecundity analysis (Eq. 1).

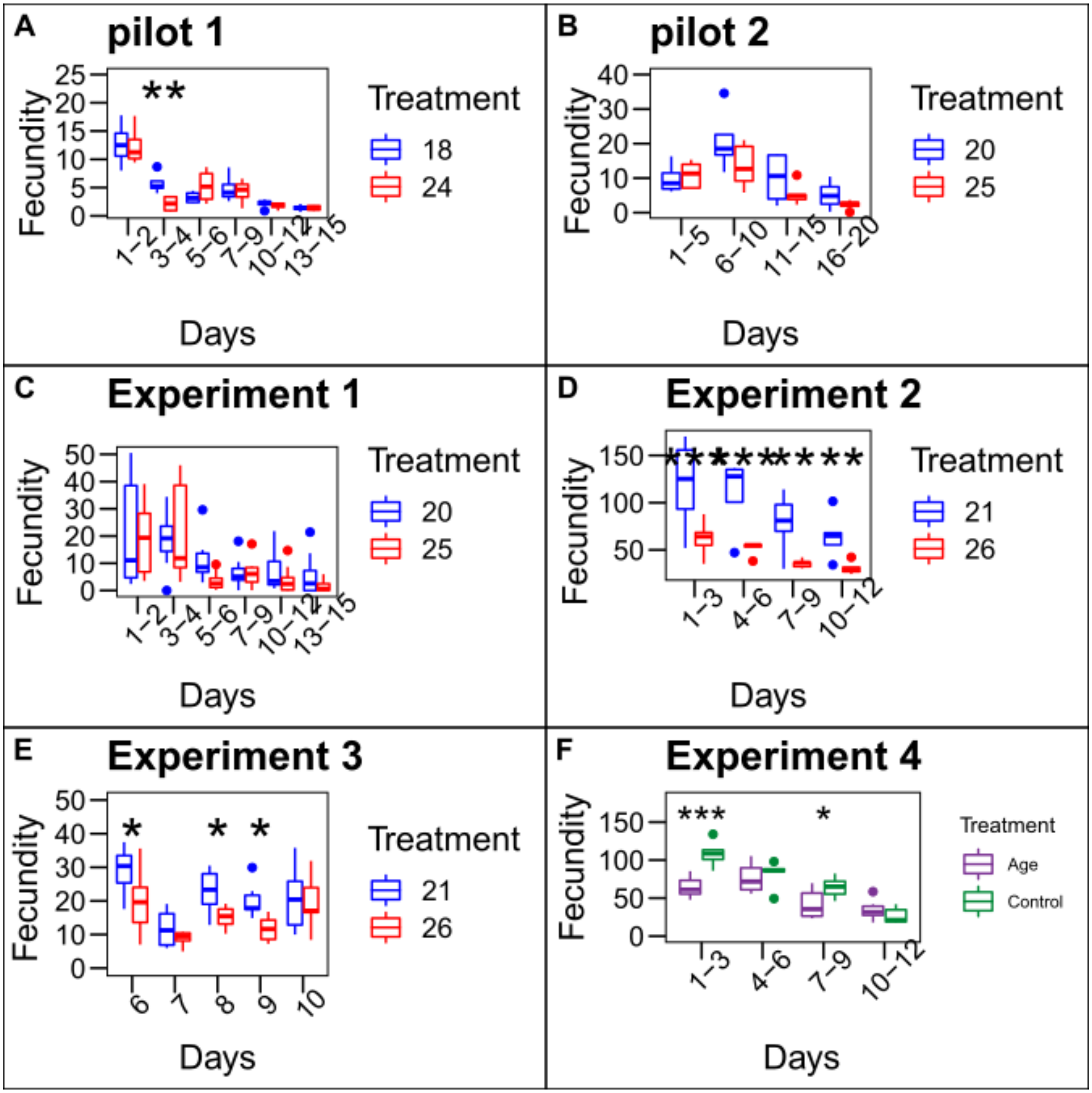


**Supplementary Figure 2. Total number of progeny (Fecundity) versus days post-mating between control and treatment across experiments.** Each boxplot represents replicate vials. As shown in Table 1, the model in Equation 2 was only significant for treatment in Experiments 2-3 (D-E). A post hoc test was done to calculate significance for each timepoint between treatment and control with significance indicated via asterisks (see Table S8).

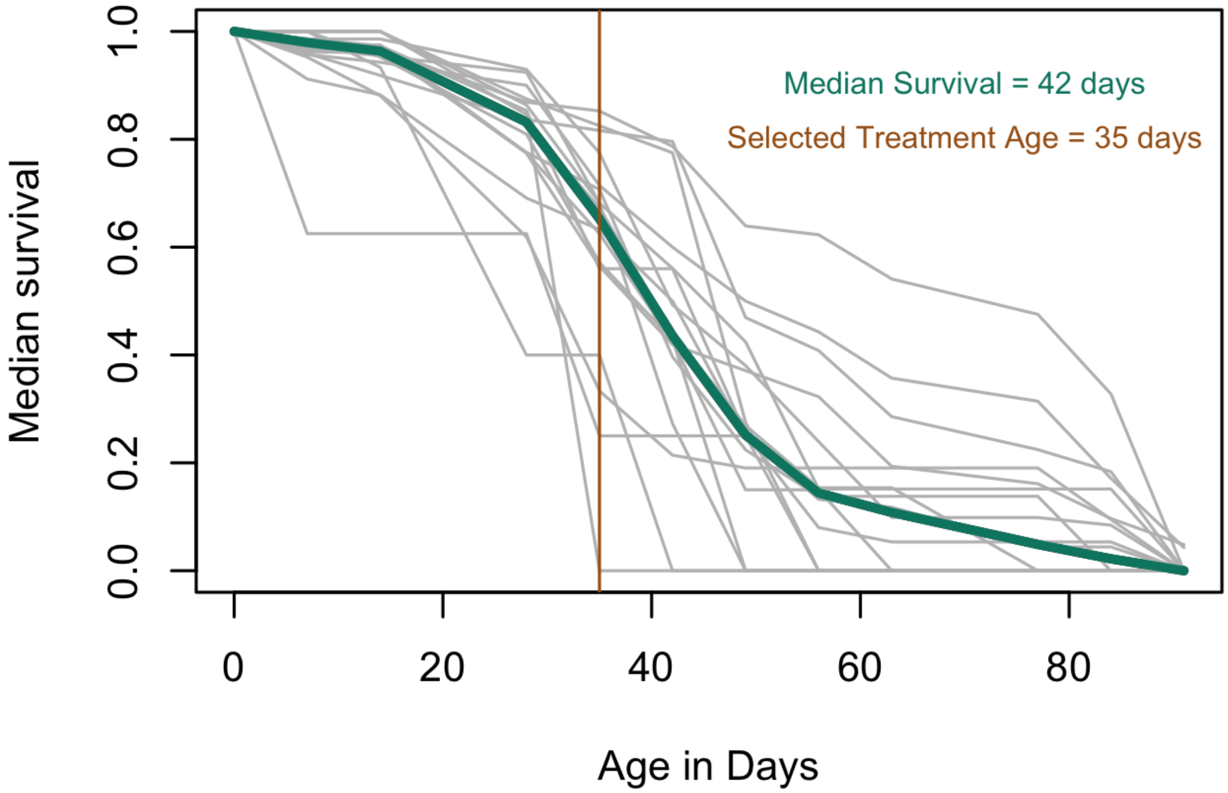


**Supplementary Figure 3. Survivorship declines with age.** Percentage of $F_1$ females remaining as a function of time. Eighteen replicate crosses were tracked until none were remaining (each shown in grey). Median survivorship for each time point across replicates is shown in dark cyan. The selected treatment for maternal age (dark orange) was the time point immediately preceding the time point where median survival fell below 50%.

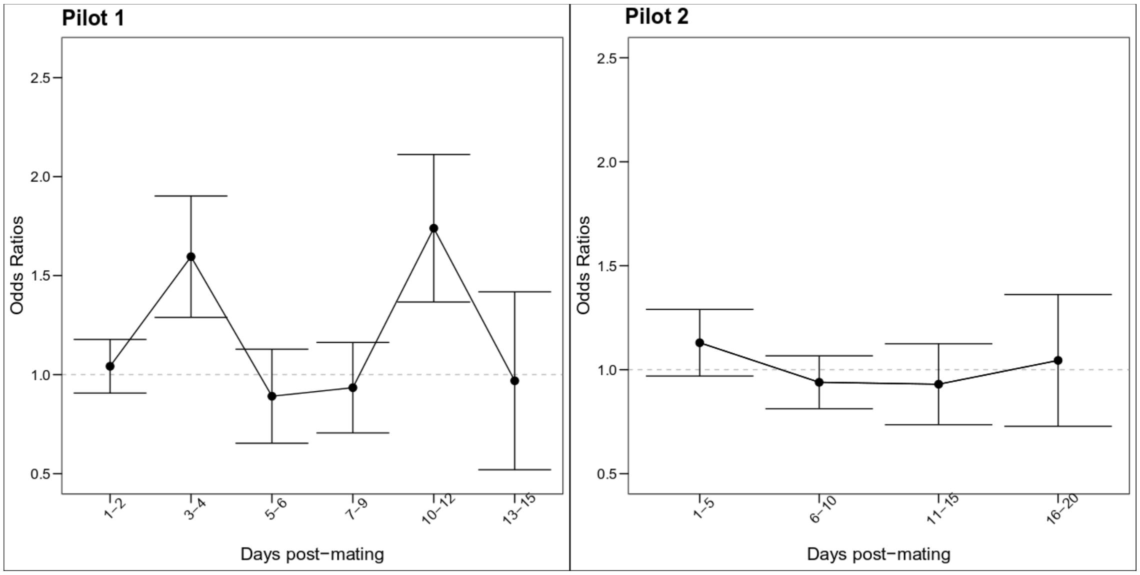


**Supplementary Figure 4. Effect of temperature on recombination rate in pilot experiments.** The first two panels describe the odds of having recombination in the treatment groups in the double mutant (*y-st*), while the last panel describes the triple mutant. The cross designs are explained in detail in the supplementary table 1. Odds were calculated from the estimates of mixed model analysis.

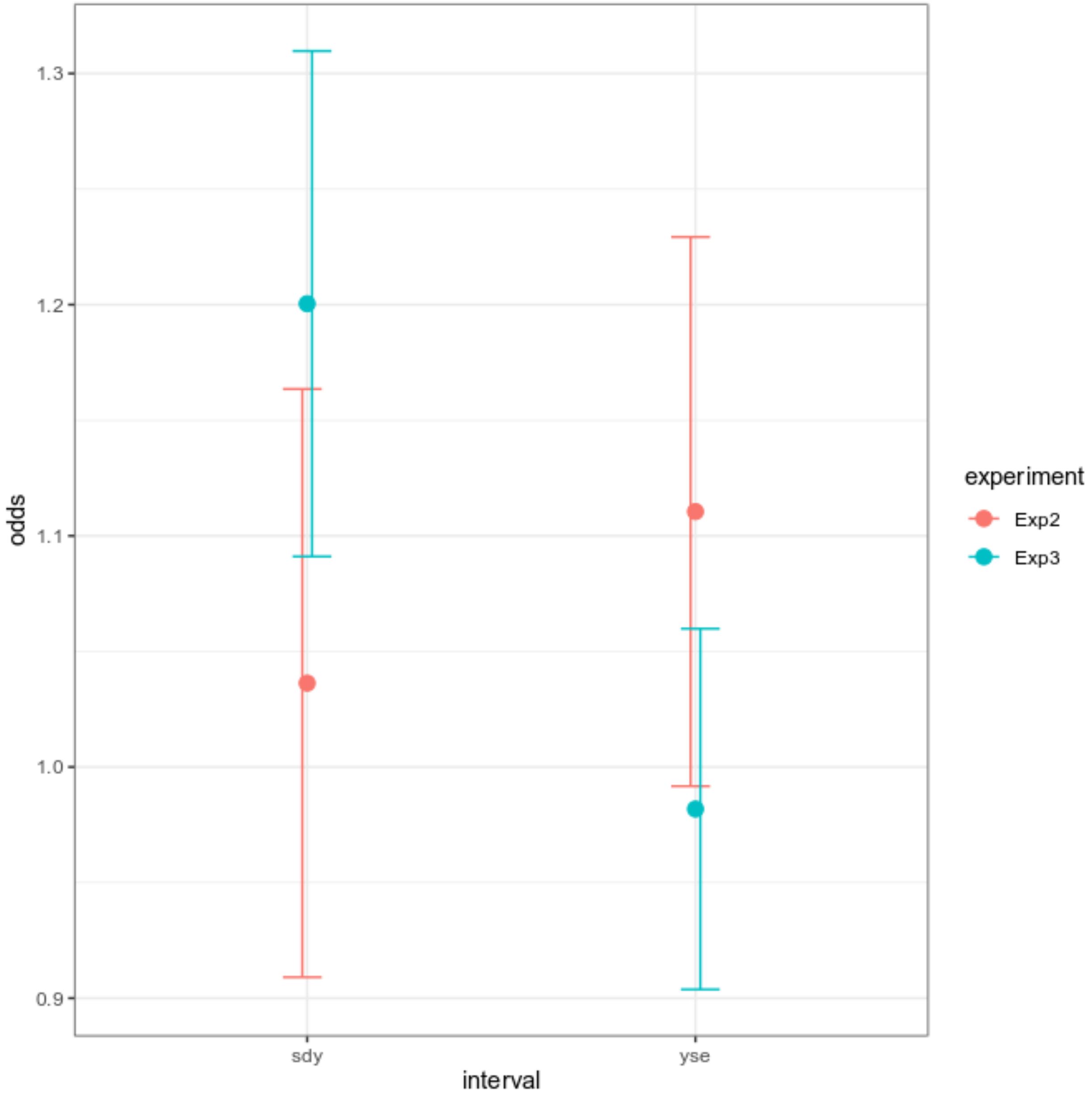


**Supplementary Figure 5. Reproducibility of results between experiments.** For days 7-9 post-mating, odds ratios were compared across experiments for the intervals *y-sd*, and *y-se*, and *y-st*. Legend indicates experiments compared by summing across multiple time points for 24H transfer frequency experiments.

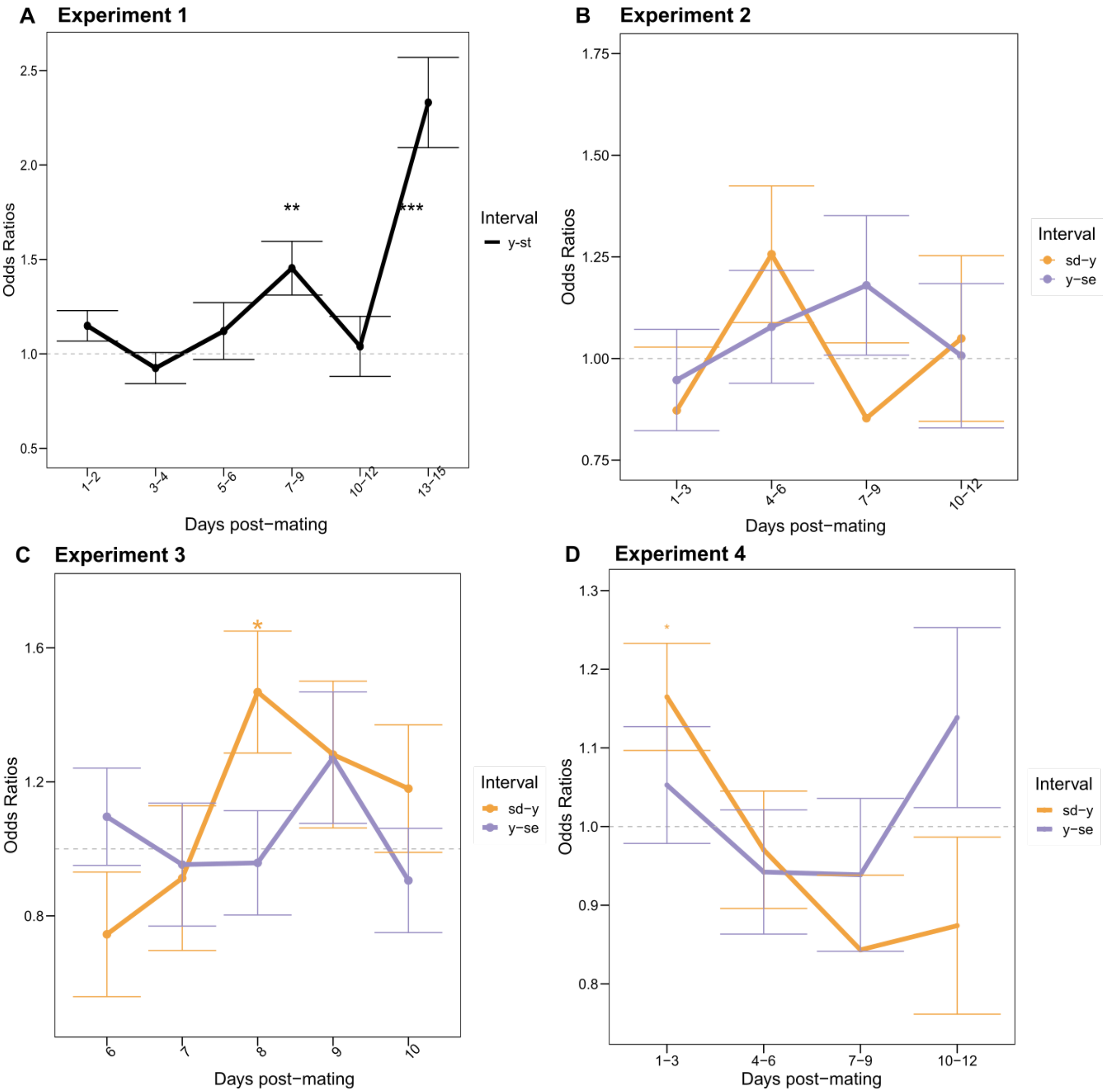


**Supplementary Figure 6. Effect of temperature on recombination rate in Experiments 1-4.** Recombination frequencies between control and treatment were compared using a fitted model using logistic regression. Exponentiating the coefficients generated the odds ratio. Odd ratios across experiments were plotted against days post-mating and indicate the odds of having a crossover in high temperature compared to control. (A) Odds ratios for the double recombination analysis in the *y-st* region, and (B-C) odds ratios for the triple recombination analysis in the intervals *sd-y* (orange) and *y-se* (purple) for Experiments 2-3. (D) Odds ratios for the triple recombination rates in response to maternal age. A *post hoc* test was done to calculate significance for each timepoint between treatment and control with significance indicated via asterisks (see Tables S5 and S6). See Table 1 and Figure 3 for additional details regarding differences in experimental design across panels

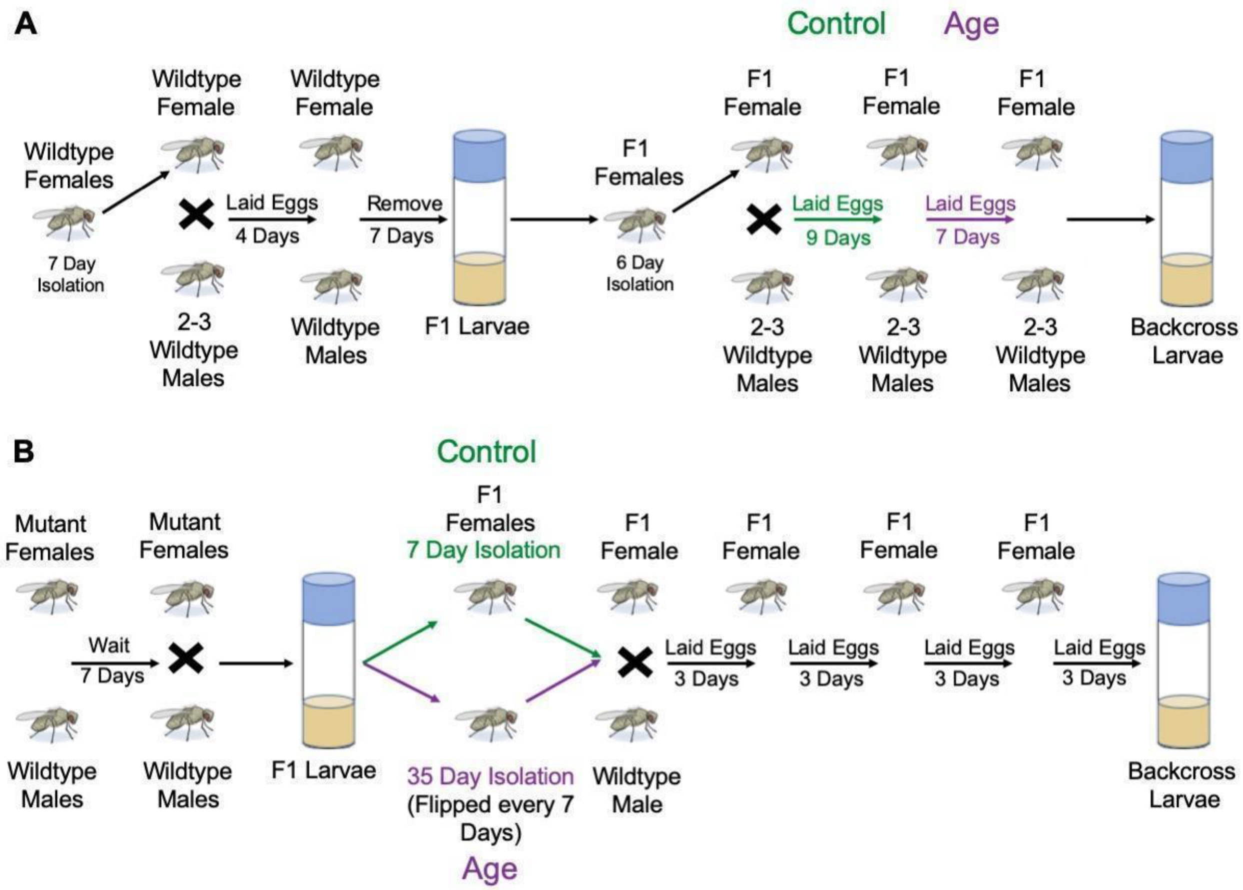


**Supplementary Figure 7. Maternal Age Experimental design.** An earlier SNP genotyping study did not find any difference in recombination rate that could be attributed to maternal age in *D. pseudoobscura*. The experimental design (A) of Manzano-Winckler *et al*. (2013), was carefully reviewed in the experimental design for this study (B). While Drosophila lay eggs continuously, artificial clutches can be made by transferring females to a fresh food vial during egg laying. A key difference in this study was that the age difference was distinct and large rather than comparing clutches approximately 1 week apart. This allowed the use of different flies for treatment and control rather than using the same flies for both.

### Supplementary Tables

**Supplementary Table 1. Summary of experimental design and results for pilot genotyping experiments to measure the impact of temperature on recombination frequency.** For each pilot experiment, a different set of temperatures, transfer frequencies and duration as well as sample sizes were used. *Note: Sample size is based only on the number of individuals targeted for recombination frequency. (e.g. only males were phenotyped). For fecundity, values are based on all progeny of both sexes for the duration of the experiment but not over the lifetime of each replicate female. For transfer frequency and duration, H denotes hours and D denotes days. ♀P-value from Eq. 1 for Treatment on Fecundity. †P-value from Eq. 2 for Treatment on Recombination Rate. Full anova tables for both analyses are in Tables S4 and S7, with post hoc tests in Tables S6 and S8. [a]Crossing scheme matches Figure S1. [ŧ]Slashes in columns two through five indicate values split by treatment and control as indicated in the temperatures column.

| | Temperatures[ŧ] | Transfer frequency /duration | # replicate crosses/treatment | Median # crosses/rep | Sample Size* | Fecundity ♀ | Recombination† |
|---|---|---|---|---|---|---|---|
| **Pilot 1[a]** | 18°C/24.5°C | 24H/15D | 4/4 | 9/8 | 1093/871 | 0.40 | 0.47 |
| **Pilot 2[a]** | 20°C/25°C | 120H/20D | 4/5 | 8/6 | 1388/953 | 0.08 | 0.98 |

**Supplementary table 2. Primers used for genotype-phenotype association.** Indel markers were designed based on the *D. pseudoobscura* reference genome 3.1. Below are the forward and reverse sequences as well as the chromosome and position. Finally, the distance to the associated gene of interest for each marker is listed.

| Gene | Name | Forward Sequence | Reverse Sequence | chr | start | end | Distance to associated gene |
|---|---|---|---|---|---|---|---|
| ***yellow*** | psy2F_M13, psy2R | ccatcggtggagaaccacta | cacgtaggctgcaattaaacg | XL_group1e | 4215723 | 4215916 | 23.2kb |
| ***vermillion*** | psv2F_M13, psv2R | atcgcctctctttcgctgag | tccgagtggcttctaacttttg | XL_group1e | 643418 | 643622 | 3.3kb |
| ***scarlet*** | DPS_Scarlet1_F_M13, DPS_Scarlet1_R | ttcactcaataatcgcagctgta | gcagtcagccaactgtgactc | XR_group8 | 2218621 | 2218787 | 21.9kb |

**Supplementary 3. SNP Genotyping recombination rate model tables.** A mixed model analysis per interval with replicate IDs as random effects and all the other parameters are fixed effects was conducted in R. The model from the GLMer was fit to an ANOVA table using chi-square with the package “car”.

| **SNP Analysis Interval 1 recombination rate** | | | |
|---|---|---|---|
| | **Chisq** | **Df** | **Pr(>Chisq)** |
| **Treatment** | 0.02 | 1 | 0.88 |
| **Day** | 3.24 | 4 | 0.52 |
| **Treatment*Day** | 5.52 | 4 | 0.24 |

| **SNP Analysis Interval 2 recombination rate** | | | |
|---|---|---|---|
| | Chisq | Df | Pr(>Chisq) |
| **Treatment** | 3.35 | 1 | 0.067 |
| **Day** | 10.94 | 4 | **0.027** |
| **Treatment*Day** | 6.22 | 4 | 0.18 |

| **SNP Analysis Interval 3 recombination rate** | | | |
|---|---|---|---|
| | **Chisq** | **Df** | **Pr(>Chisq)** |
| **Treatment** | 7.32 | 1 | **0.0068** |
| **Day** | 6.49 | 4 | 0.16 |
| **Treatment*Day** | 10.49 | 4 | **0.033** |

| **SNP Analysis Interval 4 recombination rate** | | | |
|---|---|---|---|
| | **Chisq** | **Df** | **Pr(>Chisq)** |
| **Treatment** | 4.86 | 1 | **0.027** |
| **Day** | 22.81 | 4 | **0.00014** |
| **Treatment*Day** | 11.56 | 4 | **0.021** |

| **SNP Analysis Interval 5 recombination rate** | | | |
|---|---|---|---|
| | **Chisq** | **Df** | **Pr(>Chisq)** |
| **Treatment** | 0.56 | 1 | 0.45 |
| **Day** | 10.79 | 4 | **0.028** |
| **Treatment*Day** | 5.52 | 4 | 0.23 |

**Supplementary 4. Mixed model analysis model tables for recombination rate for pilot experiments 1 and 2, and experiments 1-4.**

| Pilot Experiment 1 Recombination Rate Model | | | |
|---|---|---|---|
| | Chisq | Df | Pr(>Chisq) |
| **Treatment** | 0.52 | 1 | 0.47 |
| **Day** | 1.73 | 5 | 0.88 |
| **Treatment*Day** | 4.43 | 5 | 0.48 |
| **Pilot Experiment 2 Recombination Rate Model** | | | |
| | Chisq | Df | Pr(>Chisq) |
| **Treatment** | 0.00039 | 1 | 0.98 |
| **Day** | 2.15 | 3 | 0.54 |
| **Treatment*Day** | 1.024 | 3 | 0.79 |
| **Experiment 1 Recombination Rate Model** | | | |
| | Chisq | Df | Pr(>Chisq) |
| **Treatment** | 3.76 | 1 | **0.052** |
| **Day** | 11.225 | 5 | **0.047** |
| **Treatment*Day** | 20.14 | 5 | **0.0012** |
| **Experiment 2 sd-y Recombination Rate Model** | | | |
| | Chisq | Df | Pr(>Chisq) |
| **Treatment** | 0.086 | 1 | 0.77 |
| **Day** | 71.43 | 3 | **2.11E-15** |
| **Treatment*Day** | 1.29 | 3 | 0.73 |
| **Experiment 2 y-se Recombination Rate Model** | | | |
| | Chisq | Df | Pr(>Chisq) |
| **Treatment** | 0.01 | 1 | 0.92 |
| **Day** | 83.25 | 3 | **6.15E-18** |
| **Treatment*Day** | 2.26 | 3 | 0.52 |
| **Experiment 3 sd-y Recombination Rate Model** | | | |
| | Chisq | Df | Pr(>Chisq) |
| **Treatment** | 2.85 | 1 | 0.091 |
| **Day** | 19.02 | 4 | **0.00077** |
| **Treatment*Day** | 2.36 | 4 | 0.67 |
| **Experiment 3 y-se Recombination Rate Model** | | | |
| | Chisq | Df | Pr(>Chisq) |
| **Treatment** | 0.28 | 1 | 0.59 |
| **Day** | 14.0059 | 4 | **0.0073** |
| **Treatment*Day** | 9.048 | 4 | 0.059 |

| Experiment 4 sd-y Recombination Rate Model | | | |
|---|---|---|---|
| | Chisq | Df | Pr(>Chisq) |
| **Treatment** | 0.007 | 1 | 0.93 |
| **Day** | 4.71 | 3 | 0.19 |
| **Treatment*Day** | 9.82 | 3 | **0.02** |
| **Experiment 4 y-se Recombination Rate Model** | | | |
| | Chisq | Df | Pr(>Chisq) |
| **Treatment** | 0.0101 | 1 | 0.92 |
| **Day** | 10.08 | 3 | **0.017** |
| **Treatment*Day** | 3.18 | 3 | 0.36 |

**Supplementary 5. Post hoc table for recombination rate.** Post-hoc Analysis using Kenward – Roger model for recombination rate analysis for SNP genotyping. Means contrast using emmeans package and Kenward-Roger mode were conducted on each model in Table S3 to compare treatment and control for each time point individually.

| **SNP Analysis Interval 1 Post-hoc table** | | | | | | | |
|---|---|---|---|---|---|---|---|
| | **contrast** | **day** | **estimate** | **SE** | **df** | **z.ratio** | **p.value** |
| **1** | 23 - 18 | 1-2 | -21.28 | 512.0001 | Inf | -0.041 | 0.96 |
| **2** | 23 - 18 | 3-4 | 0.38 | 0.47 | Inf | 0.81 | 0.42 |
| **3** | 23 - 18 | 5-6 | 0.609 | 0.46 | Inf | 1.32 | 0.18 |
| **4** | 23 - 18 | 7-8 | -0.67 | 0.39 | Inf | -1.69 | 0.089 |
| **5** | 23 - 18 | 9-10 | -0.25 | 0.51 | Inf | -0.48 | 0.62 |
| **SNP Analysis Interval 2 Post-hoc table** | | | | | | | |
| | **contrast** | **day** | **estimate** | **SE** | **df** | **z.ratio** | **p.value** |
| **1** | 23 - 18 | 1-2 | -1.34 | 0.86 | Inf | -1.55 | 0.12 |
| **2** | 23 - 18 | 3-4 | 0.207 | 0.34 | Inf | 0.604 | 0.54 |
| **3** | 23 - 18 | 5-6 | 0.69 | 0.41 | Inf | 1.67 | 0.09 |
| **4** | 23 - 18 | 7-8 | 0.33 | 0.38 | Inf | 0.87 | 0.38 |
| **5** | 23 - 18 | 9-10 | 0.819218 | 0.45 | Inf | 1.82 | 0.068 |
| **SNP Analysis Interval 3 Post-hoc table** | | | | | | | |
| | **contrast** | **day** | **estimate** | **SE** | **df** | **z.ratio** | **p.value** |
| **1** | 23 - 18 | 1-2 | -0.74 | 1.12 | Inf | -0.66 | 0.51 |
| **2** | 23 - 18 | 3-4 | -0.35 | 0.55 | Inf | -0.63 | 0.52 |
| **3** | 23 - 18 | 5-6 | 1.94 | 0.62 | Inf | 3.13 | **0.0017** |
| **4** | 23 - 18 | 7-8 | 0.54 | 0.43 | Inf | 1.26 | 0.206 |
| **5** | 23 - 18 | 9-10 | 1.24 | 0.53 | Inf | 2.36 | **0.018** |
| **SNP Analysis Interval 4 Post-hoc table** | | | | | | | |
| | **contrast** | **day** | **estimate** | **SE** | **df** | **z.ratio** | **p.value** |
| **1** | 23 - 18 | 1-2 | -0.83 | 0.93 | Inf | -0.89 | 0.37 |
| **2** | 23 - 18 | 3-4 | 1.0302 | 0.41 | Inf | 2.51 | **0.012** |
| **3** | 23 - 18 | 5-6 | 0.53 | 0.61 | Inf | 0.86 | 0.38 |
| **4** | 23 - 18 | 7-8 | -0.63 | 0.52 | Inf | -1.21 | 0.22 |
| **5** | 23 - 18 | 9-10 | 1.12 | 0.46 | Inf | 2.42 | **0.015** |
| **SNP Analysis Interval 5 Post-hoc table** | | | | | | | |
| | **contrast** | **day** | **estimate** | **SE** | **df** | **z.ratio** | **p.value** |
| **1** | 23 - 18 | 1-2 | -1.12 | 1.13 | Inf | -0.98 | 0.32 |
| **2** | 23 - 18 | 3-4 | 0.89 | 0.71 | Inf | 1.25 | 0.21 |
| **3** | 23 - 18 | 5-6 | 0.43 | 0.73 | Inf | 0.58 | 0.55 |
| **4** | 23 - 18 | 7-8 | -0.35 | 0.54 | Inf | -0.64 | 0.52 |

| | | | | | | | |
|---|---|---|---|---|---|---|---|
| **5** | 23 - 18 | 9-10 | 1.39 | 0.88 | Inf | 1.58 | 0.11 |

**Supplementary 6.** Post-hoc Analysis using Kenward – Roger model for recombination rate analysis for pilot experiments 1 and 2 and experiments 1-4. Means contrast using emmeans package and Kenward-Roger mode were conducted on the model in Table S4 to compare treatment and control for each time point individually.

| Pilot Experiment 1 Recombination rate Post-hoc Analysis | | | | | | | |
|---|---|---|---|---|---|---|---|
| | contrast | day | estimate | SE | df | z.ratio | p.value |
| **1** | 24 - 18 | 1-2 | 0.041 | 0.13 | Inf | 0.3 | 0.76 |
| **2** | 24 - 18 | 3-4 | 0.46 | 0.306 | Inf | 1.52 | 0.13 |
| **3** | 24 - 18 | 5-6 | -0.11 | 0.24 | Inf | -0.48 | 0.62 |
| **4** | 24 - 18 | 7-9 | -0.068 | 0.23 | Inf | -0.29 | 0.76 |
| **5** | 24 - 18 | 10-12 | 0.55 | 0.37 | Inf | 1.48 | 0.14 |
| **6** | 24 - 18 | 13-15 | -0.031 | 0.45 | Inf | -0.07 | 0.94 |
| **Pilot Experiment 2 Recombination rate Post-hoc Analysis** | | | | | | | |
| | contrast | Day | estimate | SE | df | z.ratio | p.value |
| **1** | 25 - 20 | 1-5 | 0.121832 | 0.160132 | Inf | 0.760825 | 0.446762 |
| **2** | 25 - 20 | 6-10 | -0.06255 | 0.127125 | Inf | -0.49204 | 0.622693 |
| **3** | 25 - 20 | 11-15 | -0.07253 | 0.19471 | Inf | -0.3725 | 0.709524 |
| **4** | 25 - 20 | 16-20 | 0.043675 | 0.316876 | Inf | 0.13783 | 0.890375 |
| **Experiment 1 Recombination rate Post-hoc Analysis** | | | | | | | |
| | Contrast | day | estimate | SE | df | z.ratio | p.value |
| **1** | 25 – 20 | 1-2 | 0.14 | 0.08 | Inf | 1.72 | 0.09 |
| **2** | 25 – 20 | 3-4 | -0.08 | 0.08 | Inf | -0.95 | 0.34 |
| **3** | 25 – 20 | 5-6 | 0.11 | 0.15 | Inf | 0.76 | 0.45 |
| **4** | 25 – 20 | 7-9 | 0.37 | 0.14 | Inf | 2.63 | 8.5e-2 |
| **5** | 25 – 20 | 10-12 | 0.04 | 0.16 | Inf | 0.24 | 0.81 |
| **6** | 25 – 20 | 13-15 | 0.85 | 0.24 | Inf | 3.54 | 3.98e-4 |
| **Experiment 2 *sd-y* Recombination rate Post-hoc Analysis** | | | | | | | |
| | Contrast | Day | estimate | SE | df | z.ratio | p.value |
| **1** | 26 – 21 | 1-3 | -0.0069 | 0.11 | Inf | -0.06 | 0.95 |
| **2** | 26 – 21 | 4-6 | 0.083 | 0.12 | Inf | 0.66 | 0.51 |
| **3** | 26 – 21 | 7-9 | 0.076 | 0.12 | Inf | 0.59 | 0.55 |
| **4** | 26 – 21 | 10-12 | -0.12 | 0.15 | Inf | -0.76 | 0.45 |
| **Experiment 2 *y-se* Recombination rate Post-hoc Analysis** | | | | | | | |
| | Contrast | Day | estimate | SE | df | z.ratio | p.value |
| **1** | 26 – 21 | 1-3 | -0.12 | 0.11 | Inf | -1.11 | 0.26 |
| **2** | 26 – 21 | 4-6 | 0.026 | 0.12 | Inf | 0.22 | 0.82 |

| | | | | | | | |
|---|---|---|---|---|---|---|---|
| **3** | 26 – 21 | 7-9 | 0.059 | 0.12 | Inf | 0.48 | 0.63 |
| **4** | 26 – 21 | 10-12 | 0.123697 | 0.14189 | Inf | 0.871781 | 0.38 |
| **Experiment 3 *sd-y* Recombination rate Post-hoc Analysis** | | | | | | | |
| | Contrast | day | estimate | SE | df | z.ratio | p.value |
| **1** | 26 – 21 | 6 | 0.085 | 0.14 | Inf | 0.6 | 0.55 |
| **2** | 26 – 21 | 7 | 0.046 | 0.18 | Inf | 0.25 | 0.8 |
| **3** | 26 – 21 | 8 | 0.087 | 0.14 | Inf | 0.61 | 0.54 |
| **4** | 26 – 21 | 9 | 0.33 | 0.15 | Inf | 2.083 | **0.037** |
| **5** | 26 – 21 | 10 | 0.037 | 0.14 | Inf | 0.26 | 0.79 |
| **Experiment 3 *y-se* Recombination rate Post-hoc Analysis** | | | | | | | |
| | contrast | day | estimate | SE | df | z.ratio | p.value |
| **1** | 26 - 21 | 6 | 0.31 | 0.12 | Inf | 2.48 | 0.80 |
| **2** | 26 - 21 | 7 | -0.059 | 0.16 | Inf | -0.35 | 0.7 |
| **3** | 26 - 21 | 8 | -0.069 | 0.13 | Inf | -0.52 | 0.6 |
| **4** | 26 - 21 | 9 | 0.13 | 0.15 | Inf | 0.85 | 0.39 |
| **5** | 26 - 21 | 10 | -0.18 | 0.13 | Inf | -1.42 | 0.15 |
| **Experiment 4 *sd-y* Recombination Rate Post-hoc Analysis** | | | | | | | |
| | contrast | Day | estimate | SE | df | z.ratio | p.value |
| **1** | age -control | 1-3 | 0.15 | 0.068 | Inf | 2.24 | **0.025** |
| **2** | age -control | 4-6 | -0.029 | 0.074 | Inf | -0.4 | 0.68 |
| **3** | age -control | 7-9 | -0.17 | 0.094 | Inf | -1.79 | 0.07 |
| **4** | age -control | 10-12 | -0.13 | 0.11 | Inf | -1.19 | 0.23 |
| **Experiment 4 *y-se* Recombination Rate Post-hoc Analyis** | | | | | | | |
| | contrast | Day | estimate | SE | df | z.ratio | p.value |
| **1** | age -control | 1-3 | 0.051 | 0.074 | Inf | 0.69 | 0.48 |
| **2** | age -control | 4-6 | -0.059 | 0.078 | Inf | -0.75 | 0.45 |
| **3** | age -control | 7-9 | -0.063 | 0.097 | Inf | -0.65 | 0.51 |
| **4** | age -control | 10-12 | 0.12 | 0.11 | Inf | 1.13 | 0.25 |

**Supplementary 7. Mixed model analysis model tables of fecundity for pilots 1 and 2, and experiments 1-4.** Quasipoisson regression analysis of fecundity as a response variable and model variables of treatment and day with an interaction term was conducted in R. The resulting glm was fit to an Anova table using chi-square to generate the model table.

| **Pilot Experiment 1 Fecundity Model** | | | | | |
|---|---|---|---|---|---|
| | **Df** | **Deviance** | **Resid. Df** | **Resid. Dev** | **Pr(>Chi)** |
| **Treatment** | 1 | 0.57 | 46 | 153.83 | 0.4 |
| **Day** | 5 | 114.93 | 41 | 38.89 | **1.25E-28** |
| **Treatment*Day** | 5 | 8.67 | 36 | 30.22 | 0.059 |
| **Pilot Experiment 2 Fecundity Model** | | | | | |
| | **Df** | **Deviance** | **Resid. Df** | **Resid. Dev** | **Pr(>Chi)** |
| **Treatment** | 1 | 8.45 | 33 | 175.13 | 0.086 |
| **Day** | 3 | 84.97 | 30 | 90.15 | **1.70E-06** |
| **Treatment*Day** | 3 | 7.8 | 27 | 82.34 | 0.44 |
| **Experiment 1 Fecundity Model** | | | | | |
| | **Df** | **Deviance** | **Resid. Df** | **Resid. Dev** | **Pr(>Chi)** |
| **Treatment** | 1 | 1.847815 | 74 | 742.2707 | 0.629565 |
| **Day** | 5 | 221.266 | 69 | 521.0048 | 3.88E-05 |
| **Treatment*Day** | 5 | 16.25697 | 64 | 504.7478 | 0.842624 |
| **Experiment 2 Fecundity Model** | | | | | |
| | **Df** | **Deviance** | **Resid. Df** | **Resid. Dev** | **Pr(>Chi)** |
| **Treatment** | 1 | 340.65 | 38 | 454.16 | **5.54E-11** |
| **Day** | 3 | 174.87 | 35 | 279.28 | **6.33E-05** |
| **Treatment*Day** | 3 | 1.78 | 32 | 277.5 | 0.97 |
| **Experiment 3 Fecundity Model** | | | | | |
| | **Df** | **Deviance** | **Resid. Df** | **Resid. Dev** | **Pr(>Chi)** |
| **Treatment** | 1 | 29.34 | 58 | 206.31 | **0.00045** |

| | | | | | |
|---|---|---|---|---|---|
| **Day** | 4 | 77.84 | 54 | 128.46 | **1.39E-06** |
| **Treatment*Day** | 4 | 6.99 | 50 | 121.47 | 0.56 |
| **Experiment 4 Fecundity model** | | | | | |
| | **Df** | **Deviance** | **Resid. Df** | **Resid. Dev** | **Pr(>Chi)** |
| **Treatment** | 1 | 49.87 | 46 | 641.57 | **0.00093** |
| **Day** | 3 | 407.57 | 43 | 234 | **< 2.2e-16** |
| **Treatment:Day** | 3 | 54.17 | 40 | 179.83 | **0.0077** |

**Supplementary 8. Post hoc table for fecundity in phenotypic mutant experiments.** Means contrast using emmeans package and Kenward-Roger mode were conducted on the model in Table S1 to compare treatment and control for each time point individually.

| **Pilot Experiment 1 Fecundity Post-hoc Analysis** | | | | | | | |
|---|---|---|---|---|---|---|---|
| | contrast | day | estimate | SE | df | z.ratio | p.value |
| **1** | 24 - 18 | 1-2 | -0.024 | 0.18 | Inf | -0.13 | 0.89 |
| **2** | 24 - 18 | 3-4 | -1.0022 | 0.36 | Inf | -2.76 | **0.0056** |
| **3** | 24 - 18 | 5-6 | 0.47 | 0.32 | Inf | 1.51 | 0.13 |
| **4** | 24 - 18 | 7-9 | -0.12 | 0.29 | Inf | -0.41 | 0.68 |
| **5** | 24 - 18 | 10-12 | -0.17 | 0.45 | Inf | -0.39 | 0.69 |
| **6** | 24 - 18 | 13-15 | -0.00071 | 0.53 | Inf | -0.0013 | 0.99 |
| **Pilot Experiment 2 Fecundity Post-hoc Analysis** | | | | | | | |
| | contrast | day | estimate | SE | df | z.ratio | p.value |
| **1** | 25 - 20 | 5-Jan | 0.11 | 0.35 | Inf | 0.31 | 0.75 |
| **2** | 25 - 20 | 10-Jun | -0.43 | 0.27 | Inf | -1.55 | 0.12 |
| **3** | 25 - 20 | 15-Nov | -0.62 | 0.42 | Inf | -1.46 | 0.14 |
| **4** | 25 - 20 | 16-20 | -0.82 | 0.67 | Inf | -1.21 | 0.23 |
| **Experiment 1 Fecundity Post-hoc Analysis** | | | | | | | |
| | contrast | day | estimate | SE | df | z.ratio | p.value |
| **1** | 25 - 20 | 1-2 | -0.08 | 0.30 | Inf | -0.25 | 0.80 |
| **2** | 25 - 20 | 3-4 | -0.04 | 0.31 | Inf | -0.14 | 0.89 |
| **3** | 25 - 20 | 5-6 | -0.76 | 0.62 | Inf | -1.21 | 0.23 |
| **4** | 25 - 20 | 7-9 | -0.02 | 0.58 | Inf | -0.03 | 0.98 |
| **5** | 25 - 20 | 10-12 | -0.32 | 0.66 | Inf | -0.48 | 0.63 |
| **6** | 25 - 20 | 13-15 | -0.91 | 0.95 | Inf | -0.96 | 0.34 |
| **Experiment 2 Fecundity Post-hoc Analysis** | | | | | | | |
| | contrast | Day | estimate | SE | df | z.ratio | p.value |
| **1** | 26 - 21 | 1-3 | -0.65 | 0.19 | Inf | -3.31 | **0.00093** |
| **2** | 26 - 21 | 4-6 | -0.75 | 0.21 | Inf | -3.54 | **0.0004** |
| **3** | 26 - 21 | 7-9 | -0.79 | 0.25 | Inf | -3.11 | **0.0018** |
| **4** | 26 - 21 | 10-12 | -0.74 | 0.27 | Inf | -2.69 | **0.0069** |
| **Experiment 3 Fecundity Post-hoc Analysis** | | | | | | | |
| | contrast | day | estimate | SE | df | z.ratio | p.value |
| **1** | 26 - 21 | 6 | -0.38 | 0.18 | Inf | -2.074 | **0.038** |
| **2** | 26 - 21 | 7 | -0.28 | 0.28 | Inf | -1.032 | 0.3 |
| **3** | 26 - 21 | 8 | -0.41 | 0.21 | Inf | -1.97 | **0.048** |

| | | | | | | | |
|---|---|---|---|---|---|---|---|
| **4** | 26 - 21 | 9 | -0.54 | 0.23 | Inf | -2.35 | **0.018** |
| **5** | 26 - 21 | 10 | -0.061 | 0.19 | Inf | -0.31 | 0.75 |
| **Experiment 4 Fecundity Post-hoc Analysis** | | | | | | | |
| | contrast | Day | estimate | SE | df | z.ratio | p.value |
| **1** | Control - Age | 1-3 | 0.52 | 0.13 | Inf | 3.79 | **0.00014** |
| **2** | Control - Age | 4-6 | 0.074 | 0.13 | Inf | 0.53 | 0.59 |
| **3** | Control - Age | 7-9 | 0.43 | 0.17 | Inf | 2.47 | **0.013** |
| **4** | Control - Age | 10-12 | -0.26 | 0.22 | Inf | -1.16 | 0.24 |

**Supplementary table 9. Haplotype frequencies as in Tables 2-3, but here for pilot experiments.** Binomial test was performed to test for the deviations between paired haplotype groups in the same experimental treatment/time point that should be in equal proportions (bold values indicate significance; ***p-value<0.001, **p-value<0.01, *p-value<0.05).

| pilot experiment 1 | | | | |
|---|---|---|---|---|
| CO Class | *NCO* | | *SCO* | |
| Haplotype | *y-+* | *+-st* | *+-+* | *y-st* |
| | 451 | 469 | 504* | 585* |
| Bias Ratio | 0.96 | | **0.86** | |
| pilot experiment 2 | | | | |
| CO Class | *NCO* | | *SCO* | |
| Haplotype | *y-+* | *+-st* | *+-+* | *y-st* |
| | 607* | 535* | 600 | 599 |
| Bias Ratio | **0.88** | | 1.00 | |

| **Pilot 1** | | | | | | | |
|---|---|---|---|---|---|---|---|
| **haplotype** | treatment | 1-2 | 3-4 | 5-6 | 7-9 | 10-12 | 13-15 |
| **y+** | 18 | 92 | **36*** | 32 | 49 | 18 | 12 |
| **+st** | 18 | 114 | **58*** | 25 | 36 | 18 | 9 |
| **++** | 18 | **113*** | **41*** | 31 | 54 | 17 | 19 |
| **yst** | 18 | **146*** | **79*** | 32 | 38 | 26 | 12 |
| **y+** | 24 | 92 | 17 | 39 | 38 | 12 | 14 |
| **+st** | 24 | 97 | 20 | 40 | 25 | 21 | 6 |
| **++** | 24 | 106 | 15 | 46 | 40 | 12 | 10 |
| **yst** | 24 | 122 | 17 | 52 | 33 | 13 | 15 |

| **Pilot 2** | | | | | |
|---|---|---|---|---|---|
| **haplotype** | treatment | 1-5 | 6-10 | 11-15 | 16-20 |
| **y+** | 20 | 76 | 167 | 67 | 37 |
| **+st** | 20 | 59 | 155 | 83 | 35 |
| **++** | 20 | 91 | **135*** | 69 | 36 |
| **yst** | 20 | 73 | **175*** | 85 | 45 |
| **y+** | 25 | **104**** | 111 | 32 | 13 |
| **+st** | 25 | **55**** | 90 | 45 | 13 |
| **++** | 25 | 94 | 109 | 49 | 17 |
| **yst** | 25 | 77 | 97 | 36 | 11 |

**Supplementary table 10. Haplotype frequencies are provided as in Tables 2-3, but here broken down further by sex for Experiment 1.** Binomial test was performed to test for the deviations between paired haplotype groups in the same experimental treatment/time point that should be in equal proportions (bold values indicate significance; ***p-value<0.001, **p-value<0.01, *p-value<0.05).

| Experiment 1 | | |
|---|---|---|
| | M | F |
| **y+** | 1061 | **1327*** ** |
| **+st** | 1131 | **1089*** ** |
| **++** | **1070*** ** | **1604*** ** |
| **yst** | **1363*** ** | **1110*** ** |

| **Experiment 1** | | | | | | | | | | | | | |
|---|---|---|---|---|---|---|---|---|---|---|---|---|---|
| haplo-type | treat-ment | 1-2 | | 3-4 | | 5-6 | | 7-9 | | 10-12 | | 13-15 | |
| | | M | F | M | F | M | F | M | F | M | F | M | F |
| y+ | 20 | 178 | 196 | 146 | 172 | **77*** | 129 | **60*** | **78*** ** | 40 | **74*** ** | **57*** ** | **45*** |
| +st | 20 | 169 | 214 | 171 | 183 | **111*** | 99 | **35*** | **41*** ** | 45 | **33*** ** | **21*** ** | **24*** |
| ++ | 20 | 219 | 222 | 169 | **209**** | 102 | 102 | **49*** | **114*** ** | **18*** ** | **134*** ** | **31*** | **132*** ** |
| yst | 20 | 205 | 246 | 192 | **151**** | 113 | 125 | **75*** | **15*** ** | **84*** ** | **15*** ** | **53*** | **14*** ** |
| y+ | 25 | **184*** | 217 | 183 | 189 | 34 | 38 | 45 | **119*** ** | 43 | 36 | 14 | **34*** ** |
| +st | 25 | **236*** | 204 | 220 | 208 | 31 | 33 | 45 | **23*** ** | 35 | 24 | 12 | **3*** ** |
| ++ | 25 | 206 | 208 | **197**** | **267*** | 29 | **18*** ** | **21*** ** | **81*** ** | **20**** | **96*** ** | 9 | **21*** ** |
| yst | 25 | 216 | 245 | **256**** | **219*** | 40 | **45*** ** | **72*** ** | **19*** ** | **45**** | **16*** ** | 12 | **0*** ** |
